# *Thinopyrum intermedium* TiAP1 interacts with a chitin deacetylase from *Blumeria graminis* f. sp. *tritici* and increases the resistance to *Bgt* in wheat

**DOI:** 10.1101/2021.02.08.430348

**Authors:** Yanlin Yang, Pan Fan, Jingxia Liu, Wenjun Xie, Na Liu, Zubiao Niu, Quanquan Li, Jing Song, Qiuju Tian, Yinguang Bao, Honggang Wang, Deshun Feng

## Abstract

The biotrophic fungal pathogen *Blumeria graminis* f. sp. *tritici* (*Bgt*) is a crucial factor causing reduction of global wheat production. Wild wheat relatives, e.g. *Thinopyrum intermedium,* is one of the wild-used parents in wheat disease-resistant breeding. From *T*. *intermedium* line, we identified the aspartic acid protein gene, *TiAP1*, which involved in resistance against *Bgt*. TiAP1 is a secreted protein that accumulates in large amounts at the infection sites of powdery mildew and extends to the intercellular space. Yeast two-hybrid showed that it interacted with the chitin deacetylase (BgtCDA1) of *Bgt*. The yeast expression, purification, and *invitro* test confirmed the chitin deacetylase activity of BgtCDA1. The bombardment and VIGS mediated host-induced gene silencing showed *BgtCDA1* promotes the invasion of *Bgt*. Transcriptome analysis showed the cell wall xylan metabolism, lignin biosynthesis-related, and defence genes involved in the signal transduction were upregulated in the transgenic *TiAP1* wheat induced by *Bgt*. The TiAP1 in wheat may inactivate the deacetylation function of BgtCDA1, cause chitin oligomers expose to wheat chitin receptor, then trigger the wheat immune response to inhibit the growth and penetration of *Bgt*, and thereby enhance the tolerance of wheat to pathogens.

## Introduction

Wheat (*Triticum aestivum* L.) is one of the main crops in the world, therefore the production of wheat is closely related to food security. In recent years, pathogenic infections have seriously affected the yield of wheat. Wheat powdery mildew caused by the obligate biotrophic fungus *Blumeria graminis* f. sp. *tritici* (*Bgt*) can result in severe reductions in grain yield (Singh *et al*., 2016). When powdery mildew invades the plant, it releases proteinaceous exudates that contain the mechanical and molecular features required for the full virulence of the pathogen (Zhang *et al*., 2005). Fungal exudates function in the attachment of conidia and in the adherence of the penetrating appressoria (App) to invade plants (Deising *et al*., 2000). At the same time, the plant responds immediately to the pathogen invasion, and a number of secreted proteins, especially protease, from plant cells accumulate in the apoplastic space to carry an apoplastic battle between hosts and pathogen (Geziel, 2015; Wang et al., 2020).

In wheat, more than 66 powdery mildew resistance loci (*Pm1*-*Pm66*) located on different chromosomes of the common wheat have been named (Li *et al*., 2020a). Most of them originated from wild relative species of wheat, such as *Pm21* from *H. villosa* has been introduced into the common wheat using the translocation line T6VS·6AL (Chen *et al*., 2013). Li *et al*. (2020b) cloned *Pm41*, which was a powdery mildew resistance gene deriving from the wild emmer wheat. *Thinopyrum intermedium* had been hybridized extensively with wheat and proven a useful source of resistance to various diseases of hexaploid wheat, and several powdery mildew resistance genes, such as *Pm40*, *Pm43*, and *PmL962* (Luo et al., 2009; He et al., 2009; Shen et al., 2015), have been identified and genetic mapped. The mining and utilization of these new resistance genes are vital means to improve current wheat varieties and increase their resistance to powdery mildew.

Aspartic proteases consist of four main proteolytic enzymes that are widely present in animals, plants, yeast, microorganisms, and viruses (Rawlings and Barrett, 1995). Xia *et al*. (2004) reported that the peptide signal system of CDR1 was involved in the activation of the resistance mechanism, while Prasad *et al*. (2009, 2010) found that *OsCDR1/OsAP5* could enhance the disease resistance of rice and *Arabidopsis thaliana*. The aspartic protease gene, *AP13*, from grape could promote the salicylic acid-dependent signal transduction pathway (Guo *et al*., 2016). Moreover, Alam *et al*. (2014) found that the rice *OsAP77* gene accumulated in the extracellular space and sieve tubes to defend the pathogens. Hence, these studies have shown aspartic proteases participate in the biotic stresses of plants to enhance the resistance.

To resist pathogens, plants have developed a complex immune system including the plasma membrane receptors that recognise pathogen-related molecular patterns, such as chitin from the fungal cell walls that can trigger defence responses. Kamakura *et al*. (2002) found a new germination tube-specific gene, *CBP1*, in the rice blast fungus *Magnaporthe grisea*. This gene encodes a chitin-binding protein (CBP) having two similar chitin-binding domains at both sides of the central domain. This structure is similar to the fungus chitin deacetylase, which may play a vital role in the hydrophobic surface sensing of *M. grisea* during App differentiation. Kuroki *et al*. (2017) confirmed *CBP1* encoded a chitin deacetylase and participated in differentiation stage of App in *M. oryzae*. These results proved the chitin deacetylase activity of CBP1 is necessary for the formation of App. In addition, Yang *et al*. (2019) identified the chitinase gene *MoChia1* from *M*. *oryzae*. This gene can trigger a plant’s defence response to *M. oryzae* in rice under an inducible promoter. MoChia1 was also a functional chitinase that is required for the growth and development of *M*. *oryzae*. The MoChia1 binding to free chitin could inhibit the plant immune response, while another protein, OsTPR1, competitively binding with MoChia 1, would re-establish the immune response (Yang *et al*., 2019). Han *et al*. (2019) also found the mechanism of MoChi1 that targeted the host lectin to inhibit rice immunity and promote the colonisation. Furthermore, they also found rice protein OsMBL1 could interact with MoChi1 to enhance the resistance against *M. oryzae*. Overexpressing of *OsMBL1* can lead to the activation of defence response genes of rice and the burst of chitin-induced reactive oxygen species (ROS). Moreover, Gao *et al*. (2019) identified a secreted polysaccharide deacetylase (PDA1) from soil-borne *Verticillium dahliae*. PDA1 can promote the deacetylation of chitin oligomers, whose *N*-acetyl group contributes to the host lysine motif (LysM)-containing receptor that perceives ligand-triggered immunity and facilitates virulence. Thus, the silencing of PDA1 allows the N-acetyl group of the chitin-triggered host immunity to occur.

In a previous study, we obtained an aspartic protease gene, *TiAP1*, from trititrigia SN6306 through a comparative transcriptome analysis. Further sequence analysis showed *TiAP1* originated from *T. intermedium* (Tian *et al*., 2017). Through gene expression and virus-induced gene silencing (VIGS) analyses, we found the *TiAP1* gene might be involved in the resistance to wheat powdery mildew. In addition, subcellular localisation analysis showed that TiAP1 is of a secretory protein. During the powdery mildew invasion, TiAP1 would assemble in large numbers at the infect site. The yeast two-hybrid (Y2H) verification, luciferase (LUC) complementation imaging (LCI), and bimolecular florescent complimentary (BiFC) analysis show that TiAP1 interacts with chitin deacetylase (BgtCDA1) of *Bgt*, while BgtCDA1 could promote the invasion of *Bgt*. Therefore, we propose that TiAP1 induces the plant to resist the invasion of pathogens by interacting with BgtCDA1, which makes the chitin of the fungus inherently immune to the plant.

## Results

### The expression of *TiAP1* mediates the *Bgt*- resistance in trititrigia SN6306 to *Bgt* invasion

Since Li *et al*. (2016) analyzed the transcriptome of SN6306 and YN15, and obtained a gene that was upregulated by *Bgt* induction, which encoded an aspartic protease, while Tian *et al*. (2017) cloned the *TiAP1* gene and found that it came from *T*. *intermedium*. Here, we found that the *TiAP1* gene was expressed in SN6306 not in wheat YN15, and was upregulated by the *Bgt-*induced in the early stages of the Bgt invasion (Fig. 1A).

**Fig. 1.**
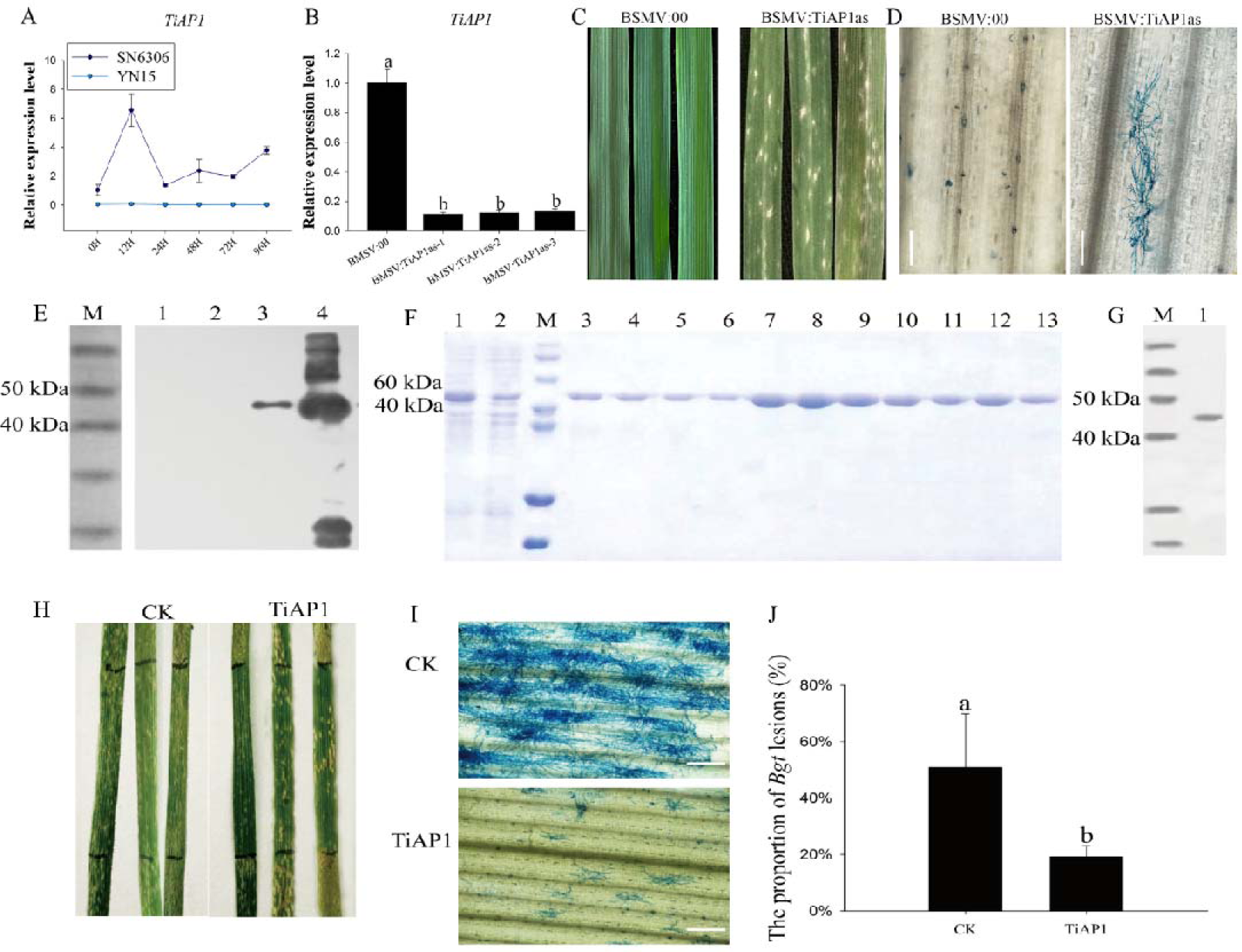
qRT-PCR and VIGS analysis of the *TiAP1* effect on the *Bgt* resistance, the TiAP1 prokaryotic expression and purification, and the reduction of the powdery mildew severity (lesions area) on the wheat seedlings by a recombination protein treatment. (A) Expression levels of *TiAP1* in YN15 and SN6306 that were induced by *Bgt*. Total RNA from the leaves of YN15 and SN6306 that were induced by *Bgt* at 0, 12, 24, 48, 72, and 96 h were extracted as the template to be detected by qRT-PCR. Data represent mean ± S.D. of *n* = 3 technical replicates. (B) Relative quantification of expression levels of *TiAP1*. Total RNA from the leaves of barley stripe mosaic virus (BSMV):00 and BSMV: TiAP1as inoculated with *E09* for ten days were extracted as the template for the detection by qRT-PCR. (C) Disease phenotype of the BSMV:00 and BSMV: TiAP1as inoculated with *E09*. Photographs were taken at 10 dpi. (D) Microscopic observation of the disease symptoms of BSMV:00 and BSMV: TiAP1as 10 dpi. Bar = 100 µm. (E) The TiAP1 protein expression detection by western blotting before and after induction. M: Pre-stained protein ladder; Lane1: Supernatant of non-induced; Lane 2: Subside of non-induced; Lane 3: Supernatant of induced; Lane 4: Subside of induced. (F) SDS-PAGE analysis of the inclusion body purification. M: Low-molecular weight protein ladder; Lane 1: Inclusions after dissolving the centrifugal supernatant; Lane 2: The effluent of supernatant was incubated with nickel iminodiacetic acid (Ni-IDA); Lane 3-6: The effluent of the imidazole elution of 50 mM; Lane 7-11: The effluent of the imidazole elution of 100 mM; Lane 12-13: The effluent of the imidazole elution of 300 mM; (G) TiAP1 quality assurance after renaturation. M: Pre-stained protein ladder; Lane 1: TiAP1 protein (1.5 µg). (H) Disease phenotype of YN15 smeared with TiAP1 and the tag protein from pET28a (CK) when inoculated with *E09* for five days. (I) Microscopic (Nikon Ni-U, Japan) analysis of the disease symptoms of YN15 smeared with TiAP1 and CK when inoculated with *E09* for five days. Bar = 100 µm. (J) The proportion of lesions in the leaf area of YN15 that was pre-treated with the recombinant TiAP1 five days before *Bgt E09* inoculation relative to the control. Control: Pre-treated with the purified recombinant label protein from the pET28a construct. Different letters above the bars indicate statistically significant differences (*P* <0.05) as obtained by one-way analysis of variance with least significance difference and Duncan’s multiple range test method.

To investigate the effect of endogenous *TiAP1* on SN6306 resistance to *Bgt*, we reduced the endogenous *TiAP1* gene levels by suppressing *TiAP1* expression using BSMV-VIGS. The *TiAP1*expression levels were significantly suppressed in the leaves of BMSV:TiAP1as SN6306 relative to the BMSV:00 negative control (Fig. 1B). When the BMSV:TiAP1as SN6306 was inoculated with *Bgt* at the two-leaf stage, the infection in BMSV:TiAP1as SN6306 was much more severe than in the BMSV:00 control at 10 days post-inoculation (dpi) (Figs. 1C,1D). This proved that the *TiAP1* gene was resistant to *Bgt* invasion and could play a role in the resistance.

### The TiAP1 protein enhances resistance to *Bgt in vitro*

Western blot analysis showed that TiAP1 was induced in the precipitation through prokaryotic expression (Figs. 1E,1F, S1). After renaturation, the protein concentration was determined to be 0.437 mg/mL by the Bradford method (Fig. 1G; Bradford, 1976). Moreover, regarding the TiAP1 expression effect on the *Bgt* invasion, the number of *Bgt* conidia on YN15 leaf after smearing TiAP1 for 5 days was significantly lower than that of YN15 with CK (pET28a). In addition, there was a small number of hyphae in the leaves smeared with TiAP1, where the hyphal length and the diseased area were less than those of the leaves smeared with CK (Figs. 1H,1I). Lesion areas of the leaves treated with TiAP1 were also significantly less than that of the control (Fig. 1J), suggesting that TiAP1 can resist the *Bgt* invasion *in vitro*.

### *TiAP1* gene transfer can elevate the resistance to *Bgt* of the recipients wheat

To investigate the function of *TiAP1*, the pMUbi-*TiAP1* vector and the screening marker *Bar* gene vector, were co-transformed into the susceptible hexaploid wheat cultivar, Bobwhite. The obtained transgenic Bobwhite with the *TiAP1* gene were subjected to PCR identification of the *Bar* and *TiAP1* genes (Figs. S2A, S2B) to obtain the positive transgenic plants. The qRT-PCR analysis showed that the expression level of the *TiAP1* gene was higher in OE1 and OE2 of SN6306 when infected by *Bgt* 2 dpi, respectively. After multiple comparisons, we found that the expression level of *TiAP1* when infected by *Bgt* in OE2-2 was significantly different from that of OE2-0 (*P* < 0.05) (Fig. 2A), indicating that the expression of *TiAP1* could be involved in the interaction with *Bgt*. In addition, we found that the overexpression of *TiAP1* gene in Bobwhite (OE1 or OE2) showed moderate resistance, while the leaf surface was still entirely green and less conidia were present at 5 dpi than the control Bobwhite (Fig. 2B). Moreover, there were a large number of conidia in Bobwhite that had germinated and formed hyphae 2 dpi using *Bgt*. A large number of conidia in the OE1-2, OE2-2 and SN6306 were retained at App of 41.2 %, 43.4 % and 83.7 %, and 16.2 %, 14.5% and 0.34 % retained at hyphae stage, respectively, while in the control Bobwhite, 28.9 % of the conidia were present in the form of App and 25.6 % of the conidia were present in the form of hyphae (Figs. 2C,2D). The HI in Bob-2 was 68.6%, while in OE1-2, OE2-2 and SN6306 it was 48.8 %, 53.5 % and 6.7 %, respectively (Fig.2F). In addition, figures 2D and 2E suggested that the transgenic *TiAP1* gene could affect the formation of *Bgt* Hau, thereby resisting the invasion of *Bgt*.

**Fig. 2.**
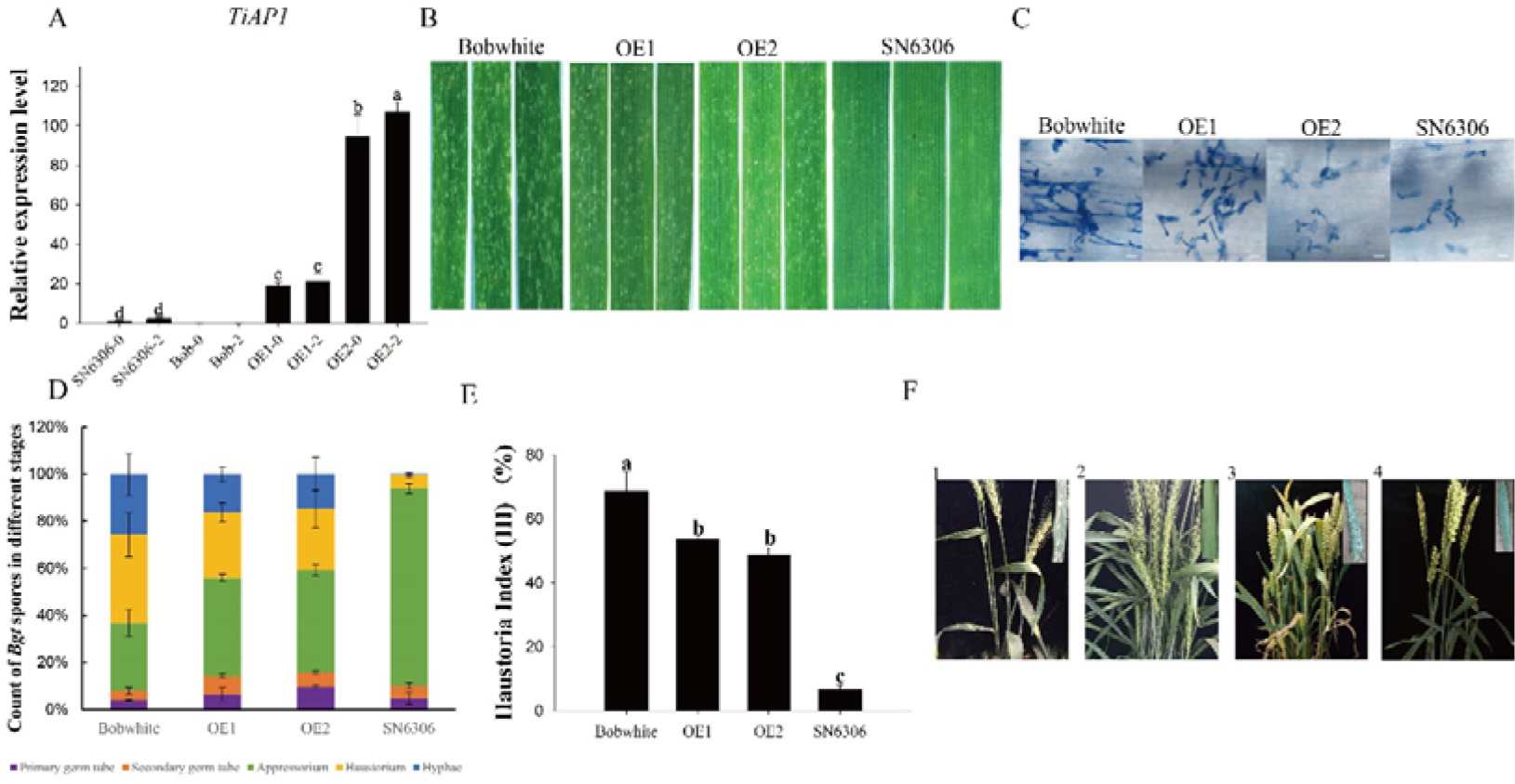
Identifying the resistance of transgenic *TiAP1* wheat. (A) Expression levels of *TiAP1* in Bobwhite and the transgenic Bobwhite with *TiAP1* when inoculated with *E09* for two days. (B) Disease phenotype of Bobwhite and the transgenic Bobwhite with *TiAP1* when inoculated with *E09* for five days. (C) Microscopic (Nikon Ni-U, Japan) analysis of the disease symptoms of Bobwhite and the transgenic Bobwhite with *TiAP1* when inoculated with *E09* for two days. Bar = 100 µm. (D) Count of *Bgt* conidia in different stages of the *Bgt* when inoculated with *E09* for two days in Bobwhite and the transgenic Bobwhite with *TiAP1*. (E) The HI statistics of the *Bgt* when inoculated with *E09* for two days in Bobwhite and the transgenic Bobwhite with *TiAP1*. Different letters above the bars indicate statistically significant differences (*P* <0.05) as obtained by one-way analysis of variance with least significance difference and Duncan’s multiple range test method. (F) Identifying the resistance of different wheat at the adult stage natural infected by powdery mildew in the glasshouse. (1) Bobwhite; (2) The transgenic Bobwhite with *TiAP1*; (3) YN15; (4) F_1_ obtained from cross breeding between YN15 and homozygous transgenic *TiAP1* Bobwhite.

Meanwhile, we crossbred between the homozygous transgenic *TiAP1* Bobwhite and YN15 to obtain the hybrid F_1_. Gene amplification and phenotype investigation confirmed the inheritance of the *TiAP1* gene and its function in resisting *Bgt* in the hybrid F_1_. A chi-square test showed that the proportion of resistant-susceptible separation in the F_2_ generation was 3:1. The subsequent F_3_ generation at the seedling stage (Figs. S2C, S3; Table S1) also double confirmed *TiAP1* endue the resistance to *Bgt* for wheat.

In the adult stage, through natural infection by powdery mildew in the glasshouse, the overexpression of *TiAP1* gene in Bobwhite showed high resistance to *Bgt* (Figs. 2F2, S4; Table S2), whereas Bobwhite (Fig. 2F1) and YN15 (Fig. 2F3) showed high susceptibility to *Bgt* on the leaves with many lesions of more than 1 mm in diameter and more hyphae. At the same time, the hybrid F_1_ between the overexpression of *TiAP1* gene in Bobwhite and YN15, showed high resistance to *Bgt*, where the diameter of the lesions was less than 1 mm, the amount of the leisons was little, the whole leaf surface was still green (Fig. 2F4). .

### TiAP1 is a secretory protein that accumulates at the infection site

To determine the localisation of TiAP1 in plant cells, we utilised the TiAP1-GFP, Δsp-TiAP1-GFP, and free GFP transient transformation in *Nicotiana benthamiana*.

Through plasmolysis methods, we found that the fluorescent signals of TiAP1-GFP were outside the cell membrane, while Δ sp-TiAP1-GFP (without signal peptide) was in the cytoplasm (Figs. 3A,3B,3C, S5). To further confirmation whether TiAP1 is a secretory protein, the mCherry-TaSYP51, which is a member of the syntaxin superfamily on plasma membrane, was co-bombarded with TiAP1-mYFP into the barley leaf epidermal cells. We found a large numbers TiAP1 and TaSYP51 proteins accumulated at the infection site of the primary and App germination tube of *Bgh*, and TiAP1 extended to the barley intercellular space (Fig. 3D; Video S1). Integrating with our observations of the *Bgt* on the leaves of *TiAP1* gene transferred wheat, we speculated that TiAP1 restrains the powdery mildew from penetration, assisting the plants in resisting the invasion of *Bgt*.

**Fig. 3.**
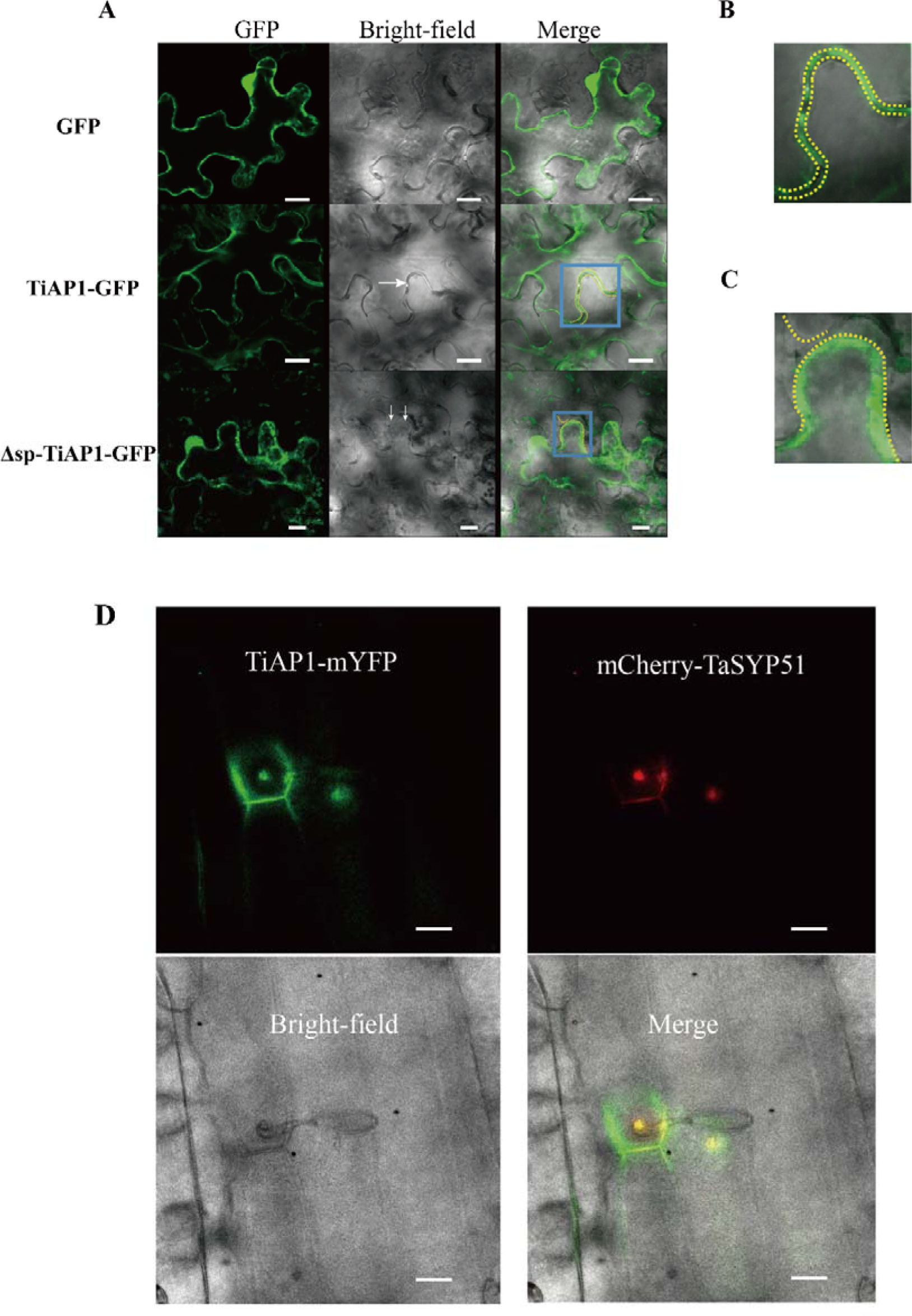
Subcellular localisation of the fusion protein through transient expression of agroinfiltrated GFP, TiAP-GFP, and Δsp-TiAP-GFP in the epidermal cells of *N*. *benthamiana* after plasmolysis, and co-transient expression of TiAP1-mYFP and mCherry-TaSYP51 in the epidermal cells of barley through bombardment. (A) Confocal laser scanning microscopy (Leica SP5-X, Germany) observation of the leaf epidermal cells revealed that Δsp-TiAP-GFP appears to be localised in the cytoplasm, while TiAP1-GFP is predominantly localised in the extracellular region, where white arrows indicate plasmolysis and the yellow dotted line shows the location of the cell membrane of the two adjacent cells. A GFP signal was used as the control. Bar = 20 µm. (B) The enlargement of the blue box in TiAP1-GFP. (C) The enlargement of the blue box in TiAP1-GFP. (C) The enlargement of the blue box in Δsp-TiAP1-GFP. (D) *B. graminis* f. sp*. hordei* induced the accumulation of TiAP1-mYFP and mCherry-TaSYP51 at the infection site. This figure is part of Video S1. Bar = 20 µm.

### The BgCDA1 interact with TiAP1

To clarify how TiAP1 can regulate wheat resistance to *Bgt*, CDS of TiAP1 without signal peptide was fused with the GAL4 DNA-binding domain sequence to generate the bait vector. We performed a Y2H screen assay using *Bgt* cDNAs as a prey library. Interestingly, one of the obtained interactors with highly similar (95.06%) to the chitin deacetylase (EPQ66796.1) of *Bgt*. Furthermore, the analysis of the cDNA revealed that the related protein contains a polysaccharide deacetylation domain. Therefore, we named this gene chitin deacetylase (*BgtCDA1*) of *Bgt* (Figs. S6A, S6B). The BgtCDA1 has an N-terminal 19-aminoacid signal peptide (http://www.cbs.dtu.dk/services/SignalP) and five conserved motifs that are required for hydrolysing the acetyl groups of the substrates (Figs. S6C, S6D). Moreover, the point-to-point verified the BgtCDA1 without signal peptide fused to the GAL4 activation domain interact with TiAP1 (Fig. 4A).

**Fig. 4.**
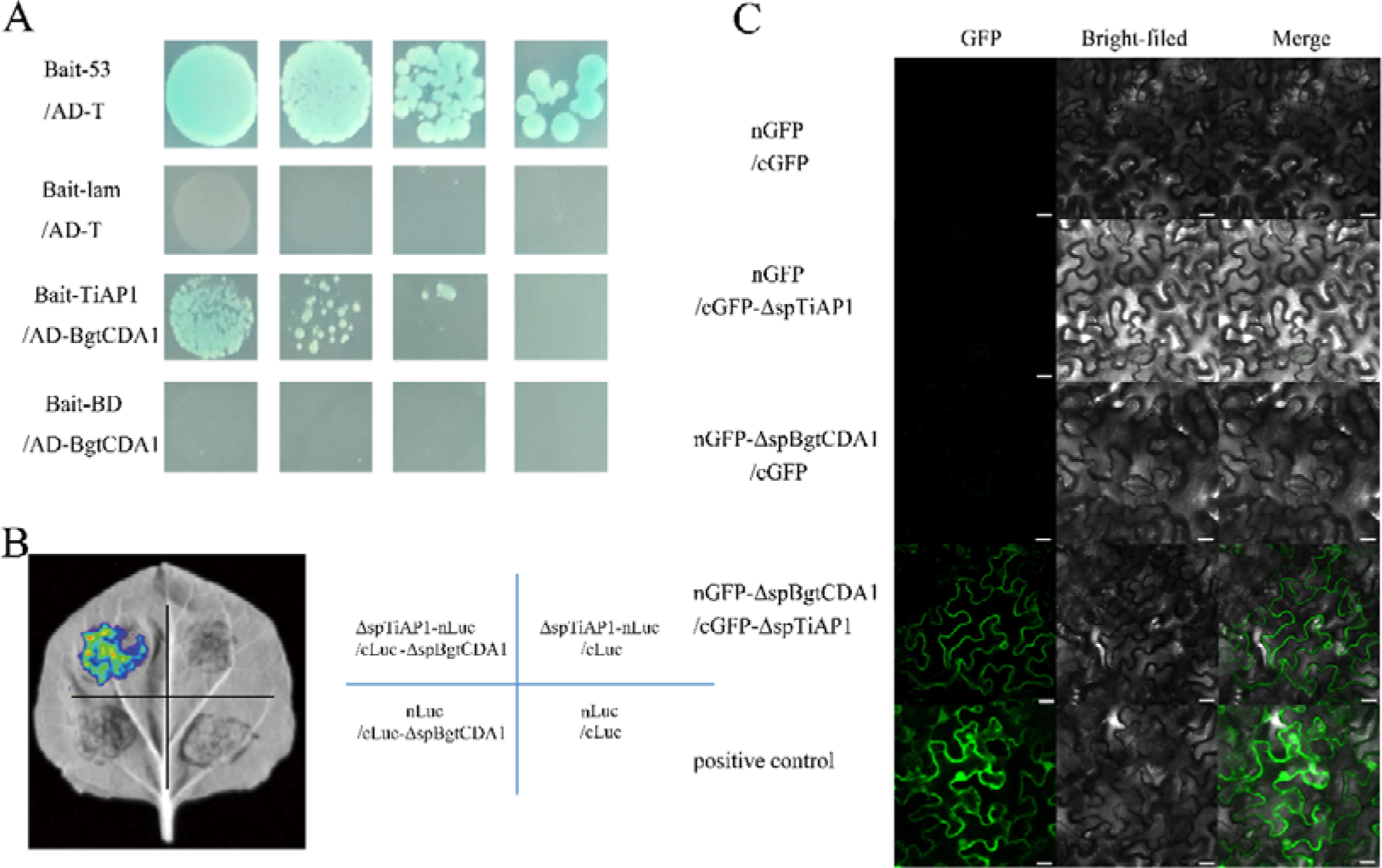
Interaction *in vitro* and *in vivo* of TiAP1 with BgtCDA1. (A) Yeast two-hybrid analysis of the interaction between TiAP1 and BgtCDA1. Yeasts expressing the interacting combinations of bait and prey were spotted on the SD-HAWL/X- α -Gal/ABA. Bait-53 and AD-T (prey) were the positive controls, while Bait-Lam and AD-T as well as Bait-BD and AD-BgtCDA1 were used as the negative controls. (B) Luciferase complementation imaging assay showing that TiAP1 interacted with and BgtCDA1. The ΔspTiAP1-nLuc and cLuc-ΔspBgtCDA1 were transiently co-expressed in *N*. *benthamiana*. spTiAP1 protein with the Co-infiltration of cLuc/nLuc-tagged ΔnLuc/cLuc-tagged spBgtCDA1 or the empty vectors were used as negative Δ controls, where the images were observed using chemiluminescence imaging two days later. (C) Bimolecular florescent complimentary assay showing TiAP1 interaction with BgtCDA1. The cGFP-Δ spTiAP1 and nGFP- ΔspBgtCDA1 were transiently co-expressed in *N*. *benthamiana*. Co-infiltrations of nGFP/cGFP-ΔspTiAP1, cGFP/nGFP- ΔspBgtCDA1, or the empty vectors group nGFP/cGFP were used as negative controls. The positive controls were nYFP-14-3-3/14-3-3-cCFP. The green florescent protein (GFP) signal was visualised using confocal microscopy (Leica SP5-X, Germany). Bar = 20 µm.

Subsequently, we furtherly confirmed the interaction through LCI, and BiFC. As expected, the LUC activity in the *N. benthamiana* leaves was high when co-injected using the ΔspTiAP1-nLuc and spBgtCDA1-cLuc, while the other three combinations were used as negative controls (Fig. 4B). The BiFC analysis also confirmed that the co-expression of nGFP-ΔspBgtCDA1 and cGFP-ΔspTiAP1 resulted in a clear GFP signal in the cytoplasm. In contrast, no visible signal was detected in any of the corresponding negative controls (Fig. 4D). Therefore, these observations indicate that there is an *in vivo* interaction between TiAP1 and BgtCDA1.

### HIGS analysis showed that the silencing of *BgtCDA1* inhibits the penetration and Hau formation of *Bgt*

To confirm the role of *BgtCDA1* in the growth and development of *Bgt*, we used a transient transformation system based on the particle bombardment of RNAi constructs (Douchkov *et al*., 2005). Through statistical analysis, we found that after the normalization treatment, the HI of the positive control, Mlo-RNAi, was 47.44%, while that of *BgtCDA1* RNAi was 55.36% (Fig. 5A, 5B). And through BSMV-VIGS, we also found that the lesions area in the leaves of BSMV: BgtCDA1as wheat infected with powdery mildew for 5 days and the hyphal density score of powdery mildew infection for 3 days were significantly lower than those of BSMV:00 wheat (Figs. 5C, 5D, 5E, 5F). This indicated that *BgtCDA1* gene silencing suppressed the growth of powdery mildew conidia and the *BgtCDA1* gene promoted the invasion of *Bgt*.

**Fig. 5.**
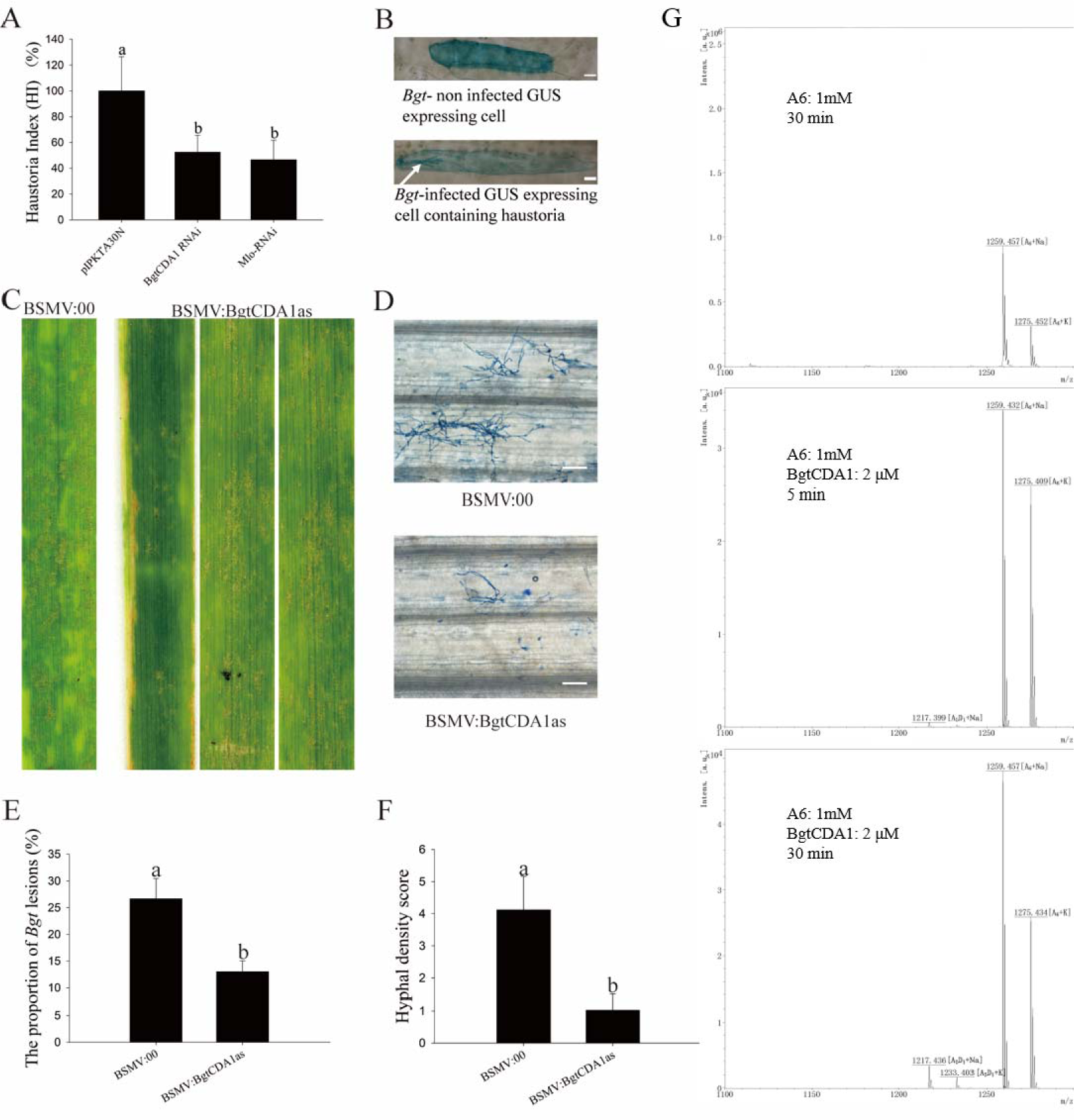
The silencing of *BgtCDA1* reduced *Bgt* infection and the BgtCDA1 mediated deacetylation of chitohexaose (A6). (A) Wheat YN15 leaves, bombarded with RNAi and GUS reporter constructs, were infected with *Bgt* and calculated for fungal haustorial index. The haustorial index was calculated as the ratio of haustoria-containing transformed cells (GUS expressing cells) divided by the total number of transformed cells. Data shown are mean values, *n* = 4, and a minimum of 150 transformed cells were counted in each repeat. Relative haustorial index was computed relative to the empty vector pIPKTA30N control of each experiment, which was set to 100%. The Mlo-RNAi was as another control. Different letters above the bars indicate statistically significant differences (*P* <0.05) as obtained by one-way analysis of variance with least significance difference and Duncan’s multiple range test method. Arrow indicates haustorium. Bar = 20 µm. (B) Wheat YN15 leaves, bombarded with RNAi and GUS reporter constructs, were infected with *Bgt.* Arrow indicates haustoria. Bar = 20 μm. (C) Disease phenotype of the BSMV:00 and BSMV: BgtCDA1as inoculated with *E09*. Photographs were taken at 5 dpi. (D) Microscopic observation of the disease symptoms of BSMV:00 and BSMV: BgtCDA1as 3 dpi using *E09*. Bar = 100 µm. (E) The proportion of lesions in the leaf area of BSMV:00 and BSMV: BgtCDA1as inoculated with *E09* five days. (F) The hyphal density score of leaf material in BSMV:00 and BSMV: BgtCDA1as 3 dpi using *E09* in microscopic observation field plots. Different letters above the bars indicate statistically significant differences (*P* <0.05) as obtained by one-way analysis of variance with least significance difference and Duncan’s multiple range test method. (G) Purified BgtCDA1 (2 uM) was incubated with 1 mM A6 for 5, 30 min as indicated. The partially deacetylated product A5D1 was detected. A, N-acetyl-D-glucosamine; D, D-glucosamine.

### The chitin deacetylase activity of BgtCDA1

We examined the chitin deacetylase activity of BgtCDA1. After Pichi yeast expression, the purified BgtCDA1-HIS (2 uM) was incubated with 1 mM Chitooctaose with six GlcNAc moieties (A6) at 37 [in 50 mM Tris-HCl buffer ( pH8.0 ), followed by detection of the enzymatic reaction products using MALDI-TOF-MS. As shown in Fig. 5G, the deacetylated intermediate products A5D1 (where A is N-acetyl-D-glucosamine and D is D-glucosamine) were detected after 30 min incubation. It showed that BgtCDA1 has the deacetylase activity.

### TiAP1 and BgtCDA1 co-regulate the expression of wheat pathogen-responsive genes

To evaluate the impact of TiAP1 and BgtCDA1 on the pathogen-responsive gene expression on a genome-wide scale, we performed RNA-Seq experiments using Bobwhite and OE2 seedlings inoculated with *E09* for two days (Fig. S7A). Genes with more than a four-fold change in expression (*P* < 0.01) were considered to be differentially expressed. Total 274 differentially expressed genes were detected in the OE2-2-vs-Bob-2 after 2 dpi, of which 116 genes were upregulated when inoculated with *E09* in the OE2-2 (Figs. S7B, S7C; Table S3). Gene ontology (GO) analysis indicated that these differentially expressed genes were enriched in the chitin catabolic process, cell wall macromolecule catabolic process, defence response to bacterium and fungus, chitinase activity, and chitin-binding (Fig. S7D; Table S4). Defence-related genes were also upregulated in OE2-2 relative to Bob-2 (Fig. 6A; Table S5), such as *traesCS1A02G410500*, *traesCS2B02G187500*, *traesCS2A02G161500*, *traesCS3B02G379200* and *traesCS1B02G440300* that encode proteins containing WRKY domains; *traesCS5A02G049600* encodes phytochrome-interacting bHLH; *traesCS7A02G201100* encodes TIFY family gene; *traesCS5D02G188600* encodes protein phosphatase 2C (PP2C); *traesCS6D02G217800* and *traesCS3A02G404400* encode ERF transcription factors; *traesCS6B02G018700* encodes LigB domain-containing protein, and may involved in betaine metabolism; *traesCS7D02G201600* and *traesCS6D02G237900* are ubiquitin E3 ligase RING-type related genes; and *traesCS6A02G266100* that encode the xyloglucan glycosyltransferase/hydrolase and may be involved in cell wall xylan metabolism. By qRT-PCR, we further analysed twelve gene which was differential expression induced of *Bgt* for two days. Among these twelve genes, the expression levels of *TraesCS1B02G440300*, *TraesCS2A02G161500*, *TraesCS2A02G199300, TraesCS3B02G379200, TraesCS5D02G188600* and *TraesCS5A02G049600* which related to plant hormone signal transduction were all upregulated in OE2-2 relative to Bob-2, beside *TraesCS5D02G188600* which was no difference in OE2-2 relative to Bob-2 (Fig. 6B). The *TraesCS3B02G259500*, *TraesCS2B02G224300*, *TraesCS6A02G266100* and *TraesCS2D02G099900* involved in cell wall anabolism, in which *TraesCS3B02G259500* and *TraesCS6A02G266100* were consistent with transcriptome were upregulated in OE2-2 relative to Bob-2, *TraesCS2B02G224300* and *TraesCS2D02G099900* were no difference in OE2-2 relative to Bob-2, but the expression in OE2-2 was higher than Bob-2, and *TraesCS6B02G018700* and *TraesCS2B02G529400* genes expression pattern were similar with *TraesCS2B02G224300* and *TraesCS2D02G099900* (Fig. 6B). These quantitative results were slightly different to those at the transcriptome, possibly because of the differences between individuals. In summary, the upregulated expression of these genes provided favourable evidence to understand the disease resistance mechanism of the *TiAP1* gene.

**Fig. 6.**
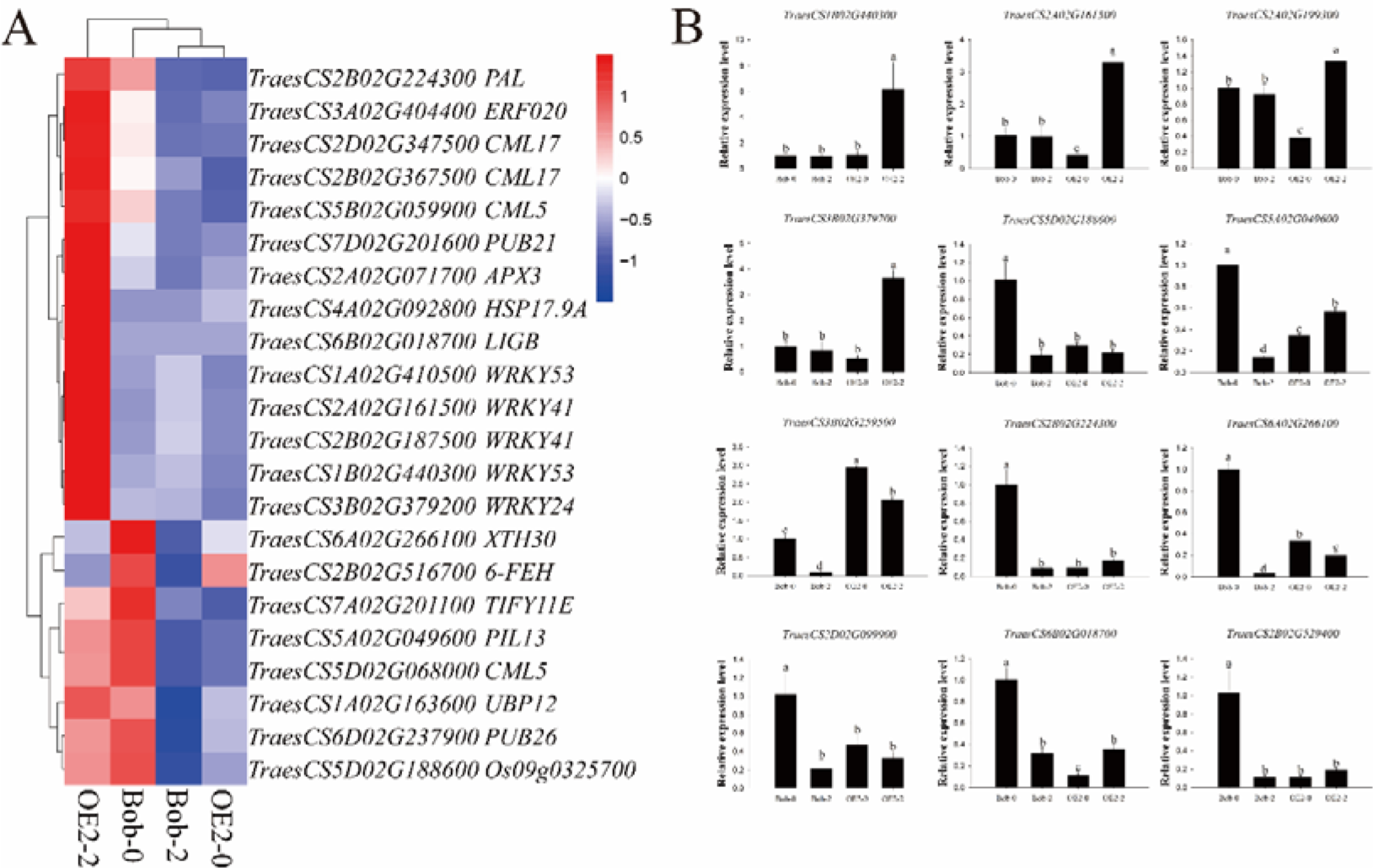
The transcriptome and qRT-PCR analysis of the transgenic wheat with *TiAP1*. (A) Heat map of selected pathogen-responsive genes in Bobwhite without and with induction of *Bgt* for two days (Bob-0 and Bob-2, respectively), and overexpression line OE2 as the same treatment (OE2-0 and OE2-2, respectively). (B) qRT-PCR analysis of the differential expression gene *TraesCS1B02G440300*, *TraesCS2A02G161500*, *TraesCS2A02G199300*, *TraesCS3B02G379200*, *TraesCS5D02G188600*, *TraesCS2B02G224300*, *TraesCS5A02G049600*, *TraesCS6A02G266100*, *TraesCS3B02G259500*, *TraesCS2D02G099900*, *TraesCS6B02G018700* and *TraesCS2B02G529400* of selected induced of *Bgt* for two days.

## Discussion

Aspartic proteases play a vital regulatory role in plant growth and may be involved in the senescence process (Chen *et al*., 2015), plant reproduction (Niu *et al*., 2013), programmed cell death (Chen and Foolad, 1997), resistance to pathogens (Prasad *et al*., 2010), response to stress (Guo *et al*., 2013) and endoproteases in plants (Rawlings *et al*., 2016). If the endoproteases encoding genes were deleted or inhibited, the plant’s sensitivity to pathogens increases (Jashni *et al*., 2015; Díaz *et al*., 2018). In the early stage, our laboratory cloned an aspartic protease (Tian et al., 2017). Through multiple sequence comparisons, we found that the TiAP1 gene is 98% similar to scaffold 208 assembled from the *T*. *intermedium* genome (https://jgi.doe.gov/), and Yang and Feng (2020) analysed the aspartic protease gene family and their response under powdery mildew stress in wheat. Among those family genes, the AP gene which most similar to *TiAP1*, TRIAE_CS42_5DS_TGACv1_456657_AA1475640.1, were located on the chromosome 5DS with 70% identity. The qRT-PCR analysis showed that the *TiAP1*gene was rapidly expressed under the induction of *Bgt* in SN6306 with no expression in YN15 (Fig. 1A). *TiAP1* expression levels were significantly suppressed with more hyphae piles appearing in the leaves of BMSV:TiAP1as SN6306 (Figs. 1B, 1C, 1D) as well as lesion areas on the leaves that were smeared with the TiAP1 protein level when compared to the control (Figs. 1H–1J). These results indicated that *TiAP1* might resist the invasion of powdery mildew.

Furthermore, the identification of the *Bgt* resistance phenotype at the seedling stage showed that the overexpression of *TiAP1* gene in wheat was moderately resistant to powdery mildew (Fig. 2B), while the life cycle of *Bgt* in the two-day-infected leaves of the overexpression of *TiAP1* gene in Bobwhite was primarily retained in the App stage. In contrast, the conidia on the Bobwhite leaves were mainly in Hau stage. Hau was wrapped in the extrahaustorial membrane (EHM), which was usually used for signal exchange and nutrient absorption (Panstruga, 2013), where the development of the Hau reflected the growth and metabolism of the entire pathogen to a certain extent. It was a vital trait for evaluating host cell resistance (Bracket, 1968).

The HI of *Bgt* in Bobwhite was significantly higher than that of the transgenic *TiAP1* gene lines, indicating that overexpression of *TiAP1* gene in Bobwhite were resistant to the *Bgt* invasion (Fig. 2E). This resistance may be because the transgenic wheat inhibited the penetration of the powdery mildew, thereby affecting their infection rate. Subsequently, we found that the transgenic *TiAP1* Bobwhite and the F_1_ generation showed high resistance to powdery mildew in the adult-stage (Figs. 2F2, 2F4, S4). Many studies refer to this trait, which showed less significant resistance to the disease at the seedling stage than that of the adult stage (Hautea *et al*., 1987; Griffey and Das, 1994), or of a chronic disease resistance type (Shaner, 1973). Moreover, Shaner (1973) found condia of Konx, a typical slow-white powder variety, could not successfully invade. Therefore, we speculated that the transgenic wheat line with the *TiAP1* gene might be used as a chronic powdery mildew-resistant variety to prevent the invasion of *Bgt*. Syntaxins were members of the SNARE superfamily of proteins that mediated membrane fusion events (Colline *et al*., 2003). Since TiAP1 was a secreted protein (Figs. 3A, 3B, 3C) which expressed in large quantities at intercellular space of the infection site (Fig. 3D, Video S1), we speculate that syntaxin TaSYP51 may help to transport the TiAP1 protein to the outside of the cell, where the detailed mechanisms need to be further elucidated.

Through the Y2H, LCI, and BiFC *in vivo* verification, we determined TiAP1 interacts with BgtCDA1 (Fig. 4). Subsequently, we found that when the *BgtCDA1* gene was silenced, the HI, lesions area and hyphal density score were significantly reduced (Fig. 5). Similarly, Zhao *et al*. (2010) found that chitin deacetylase was secreted when it interacted with other substances. The chitin deacetylase of *C. lindemuthianum* was secreted during the process of fungal hyphae invading plants to modify the chitin that can be recognised by the plant’s resistance system (Tsigos and Bouriotis, 1995). Geoghegan and Gurr (2017) also found in *M. oryzae* that CDA1 and CDA4 were necessary for chitin deacetylation in the cell wall of the mycelium. During infection, the ability of the fungal cells to convert chitin to chitosan was critical for masking the cell wall chitin from being recognised by the host chitinases, thereby promoting the fungal attack (Upadhya *et al*., 2018). Chitin oligomers can also act as ligands that were recognised by LysM receptors on the cell surface and triggered pathogen-associated molecular pattern (PAMP)-triggered immunity (PTI) (Hurlburt *et al*., 2018; Gao *et al*., 2019). We found the silencing of *BgtCDA1* can inhibit the invasion of *Bgt* and speculated that the interaction between BgtCDA1 and TiAP1 made BgtCDA1 be degraded by TiAP1 which inhibit the deacetylation function of BgtCDA1. Moreover, the growth and penetration of *Bgt* was inhibited and the chitinase of the wheat was secreted to release the chitin oligomers, which were recognised by the wheat chitin receptor, such as LysM. This led to PTI and caused the transgenic wheat with the *TiAP1* gene to be resistant to powdery mildew (Fig. 7). When the pathogen invades the host, an extremely complex and precise arms race of ‘attack and defence’ occurs in the apoplastic space. Xia *et al*. (2020) found that soybean could secreted an apoplastic aspartic protease, GmAP5, to bind and degrade a pathogen-secreted apoplastic endoglucanase PsXEG1 to destroy its virulence in *Phytophthora sojae* invasion, and proposed a ‘multi-layered immune model’ of the plants against pathogens in the extracellular region. Therefore, this study revealed the complexity of the interaction between pathogens and hosts as well as provided a theoretical basis to further study the role of TiAP1 in resistance mechanisms.

**Fig. 7.**
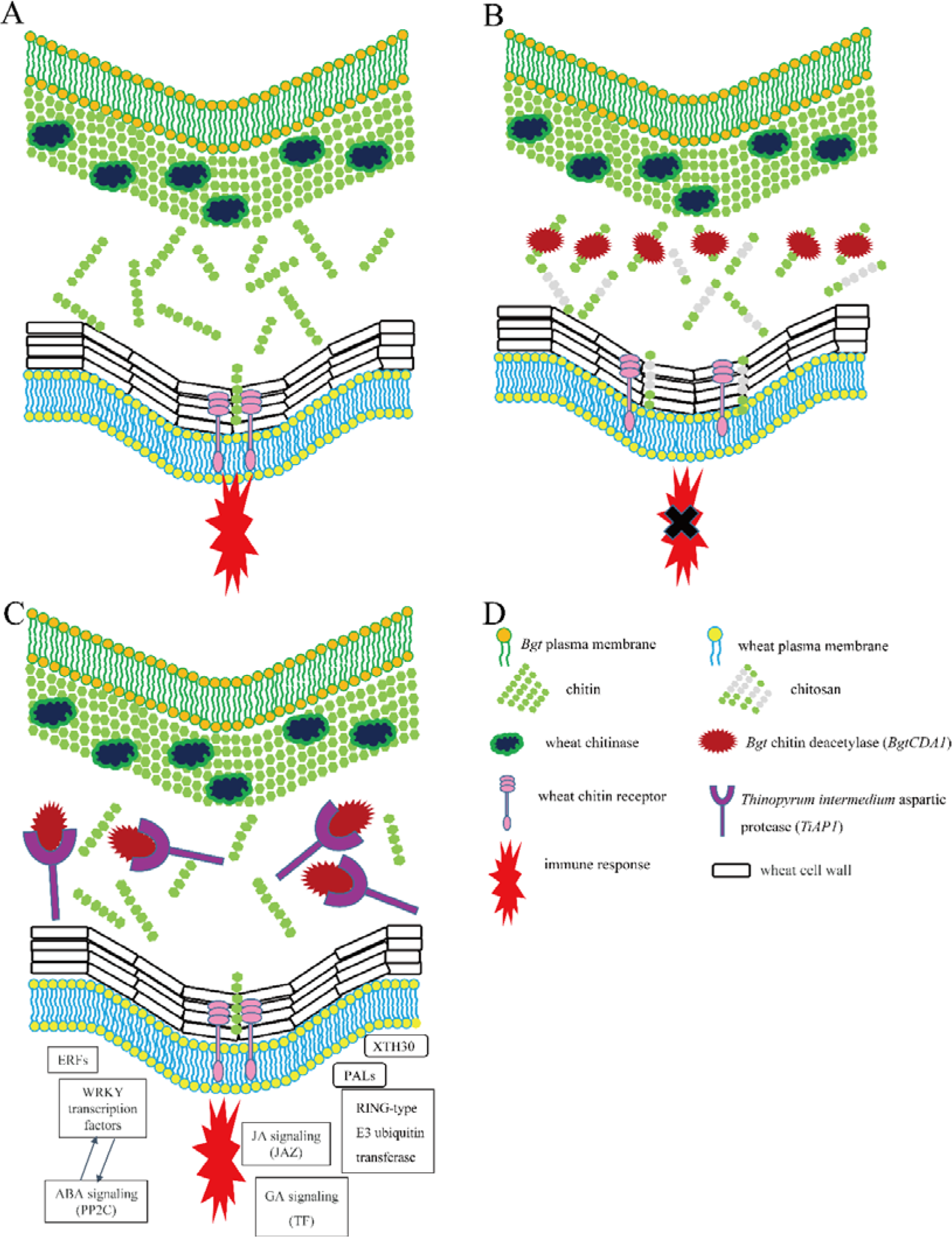
A proposed model of the TiAP1 interaction with chitin deacetylase to increase the resistance to *Bgt* in wheat. (A) The *Bgt* cell walls were targeted by wheat-secreted chitinases to release the chitin fragments that can further activate the host immune system. (B) *Bgt* would secrete BgtCDA1 to convert the cell wall chitin into chitosan, while the host invasion may protect the hyphae of the pathogenic fungi from being hydrolysed by the extracellular plant chitinases, thereby protecting the integrity of the fungal cell wall and inhibiting the resistance response mechanism of the plant. (C) Interaction between TiAP1 and BgtCDA1 in the transgenic *TiAP1* gene wheat, causing BgtCDA1 to inactivate its deacetylation function, therefore the chitin in fungal cell walls would be targeted by the wheat-secreted chitinases that liberate chitin the fragments that can further activate the host immune system. When the transgenic *TiAP1* wheat was infected by powdery mildew, the hormone signals related gene expression *in vivo* would interact to activate plant disease resistance and upregulate the expression of phenylalanine ammonia lyases in the lignin biosynthesis and XTH30 in the xylan metabolism of the transgenic *TiAP1* wheat. Hence, it enhances the ability of the plant cell walls to resist the penetration by pathogens.

When chitin receptors recognise the chitin, the plant can active defence genes and release some metabolites that initiate the signal transduction in plant cells to trigger an immune response. Many defence-related genes were upregulated in OE2-2 relative to Bob-2 (Fig. 6A; Table S5). WRKY TFs are key nodes in abscisic acid (ABA)-responsive signaling networks (Rushton et al., 2012) and Zhang et al. (2015) found the OsWRKY24, -53, and -70 are involved in gibberellic acid. Therefore, we speculate that these genes may be involved in the ABA or gibberellic acid (GA) signaling pathways. Phytochrome-interacting bHLH. These proteins are known to play important roles in red light-mediated (phyA and/or phyB-dependent) signal transduction pathways at immediate positions downstream of the photoreceptor in *A. thaliana* (Nakamura et al., 2007). In *A*. *thaliana*, the TIFY family may be involved in jasmonic acid (JA) pathway (Chung and Howe, 2009). PP2Cs are vitally involved in ABA signaling (Park et al., 2009). The ERF transcription factor family are involved in ethylene-activated signaling pathway (Xu et al., 2008). The metabolites include ABA, ethylene, JA, GA, and ubiquitination that caused the plants to respond to the invasion of pathogens (Fig. 7; Table S3). Moreover, RNA-Seq and qRT-PCR analysis revealed that there was a cell wall xylan metabolism gene, *traesCS6A02G266100*, whose expression in Bobwhite was sharply reduced after being induced by powdery mildew, while the gene in OE2-2 was significantly upregulated relative to that of Bob-2 (Fig. 6).

Since xylan was a type of hemicellulose that was closely related to functional proteins in the cell wall, it had a great influence on the ductility, pore elasticity, flow, and rheological properties of plant cells during growth (Zhou *et al*., 2017). Therefore, we infer that the penetration during the powdery mildew infection of the transgenic wheat with the *TiAP1* gene was blocked. We also found that the *TraesCS2B02G224300* gene encoded a phenylalanine ammonia-lyase (PAL), which was upregulated and significant difference in OE2-2 relative to Bob-2 in transcriptome, (Fig. 6A; Table S5) and in qRT-PCR the expression of *TraesCS2B02G224300* in OE2-2 was twice higher than Bob-2 (Fig. 6B) and the PAL was pivotal for the enzymes related to the biosynthesis of lignin, salicylic acid, and other phenylalanine-derived metabolites (You *et al*., 2020). As a main component of the plant cell wall, lignin contributes to the formation of the physical and chemical barriers in the plant immunity. Zhou *et al*. (2018) also found that *OsPAL1*-mediated resistance to rice blast might be due to the accumulation of lignin, while the broad-spectrum blast-resistant rice varieties *Pigm-* and *PIBP1-*mediated resistance might be related to lignin content (Zhai *et al*., 2019). Similarly, the overexpression of *OsPAL8* in rice can lead to increased lignin content and enhanced brown plant hopper resistance (He *et al*., 2020). Thus, these results demonstrate that the transgenic wheat with the *TiAP1* gene could enhance resistance to *Bgt*.

In summary, we speculate the fungal chitin deacetylase BgtCDA1 can block the PAMP processes that the fungal derived chitin oligoes are recognized by wheat chitin receptor protein and then trigger an immune response, while TiAP1 competitively interacting with BgtCDA1 is the causal reason to re-establish resistance to fungal pathogen *Bgt*. Moreover, the overexpression of *TiAP1* upregulates the cell wall-related genes in transgenic *TiAP1* wheat thereby blocks the penetration of powdery mildew and enhances the tolerance to pathogens. Furtheremore, we conclude many other plants may share similar mechanism and this mechanism can be used for future genetic improvement of crops. However, the mechanism of BgtCDA1 degraded by TiAP1, and the resulting detailed disease resistance pathway remains to be studied.

## Materials and methods

### Plant material and pathogen infection

The trititrigia SN6306 was highly resistant to the powdery mildew and obtained by hybridisation between the highly susceptible to powdery mildew wheat (*Triticum aestivum* L.) cultivar Yannong 15 (YN15) and *T. intermedium* (Li *et al*., 2016). Wheat Bobwhite were highly susceptible to the powdery mildew. The transgenic overexpression *TiAP1* in Bobwhite line 1 and 2 were named as OE1 and OE2, respectively. The untreated with *Bgt* wheat lines Bobwhite, OE1 and OE2 were named as Bob-0, OE1-0 and OE2-0, respectively. The Bobwhite, OE1 and OE2 wheat inoculated with *E09* for two days were named as Bob-2, OE1-2 and OE2-2, respectively. The seeds of these materials above and *N*. *benthamiana* were sown in a mixed soil (loess, matrix, and vermiculite of 1:1:1, *v*/*v*/*v*) at 25°C, 70% humidity, and a long-term photoperiod of a 16 h/8 h light/dark cycle in a glasshouse with a light intensity of 600 mmol m^-2^s^-1^. The virulent isolate *E09* of the wheat powdery mildew fungus (*Bgt*) was propagated on the wheat variety YN15 via a weekly inoculum transfer. The barley (*Hordeum vulgare* L.) line Ingrid *ror2* (Collins *et al*., 2003) was highly susceptible to *Blumeria graminis* f. sp. *hordei* (*Bgh*) and used for the localisation and expression profiling of *TiAP1*. The *Bgh* isolate B103 was propagated on barley P-10 via a weekly inoculum transfer, while the *Escherichia coli* (*E*. *coli*) strain BL21 (DE3) was used for the prokaryotic expression of TiAP1.

### VIGS, barley stripe mosaic virus (BSMV), and host-induced gene silencing (HIGS)

VIGS analysis was conducted according to Yuan *et al*. (2011) to insert the conserved sequence of the target *TiAP1* and *BgtCDA1* gene (316 bp and 300 bp, <u>respectively</u>) into the BSMV vector to create BSMV:TiAP1as and BSMV:BgtCDA1as and transform the *Agrobacterium*. The detailed method was described in Methods S1. Ten days after inoculation with BSMV, 6 leaves were harvested to test the expression level of *TiAP1* and *BgtCDA1* gene, while the other 6 leaves were inoculated with *Bgt* race *E09* and were observed and used to evaluate the resistance to the powdery mildew using a biological microscope (Nikon Ni-U, Japan) after the Coomassie Brilliant Blue staining and the statistics of hyphal density score refer to the method of Norriss et al. (2007) and three biological replicates.

*BgtCDA1* gene silencing in *Bgt* was performed using the HIGS method as described by Nowara *et al*. (2010). The RNA interference (RNAi) construct BgtCDA1 RNAi and a β-glucuronidase (GUS) reporter gene construct were co-transformed into the leaf epidermal cells of the wheat YN15 that grew for seven days using particle bombardment. The fungal haustorial formation was examined for the transformed (GUS expressing) blue cells using a biological microscope (Nicon Ni-U, Japan). The empty vectors pIPKTA30N, and the Mlo-RNAi (pIPKTA36) constructs, were used as negative and positive controls, respectively and four biological replicates were conducted per analysis with three technical replicates. The detailed method was described in Methods S1 according to Ahmed (2015).

### Constructing the TiAP1 prokaryotic expression plasmid and the protein expression

To improve the expression of *TiAP1*, the gene sequence was optimised according to *E. coli* bias, and constructed into the pET28a vector (containing 6xHis tags). Recombinant plasmids were transformed into BL21 (DE3) chemically competent cells where a single-colony transformant was cultured and inoculated at 15 °C overnight induced expression with 0.25 mM isopropyl β- d-1-thiogalactopyranoside (IPTG). The cells were harvested and subjected to a sodium dodecyl sulphate-polyacrylamide gel electrophoresis (SDS-PAGE) to detect the protein expression. Then, the recombinated TiAP1 was purified according to the method described in Methods S1.

### Determination of the antifungal activity of TiAP1

To confirm the effect of TiAP1 protein on *Bgt*, we used the renatured TiAP1 and pET28a tag protein to add 0.025% Tween-20 to smear the leaves of YN15 and inoculate with *E09*. After five days, the infectivity of *Bgt* was observed and the leaves were stained with the Coomassie Brilliant Blue stain, as described previously (Göllner *et al*., 2008) and three biological replicates were conducted per analysis with three technical replicates. Lesion areas on the leaf surface were calculated according to Goodwin and Hsiang (2010) to analyse the pixels of the *Bgt*infection area relative to the normal leaf area.

### Constructing the particle bombardment genetic transformation vector to transform wheat and identify the disease resistance phenotype

The coding sequence (CDS) of *TiAP1* (KJ513672) was cloned into the *Ubi*-*gene*-Tnos vector with a ubiquitin (Ubi) promoter, yielding a recombinant pMUbi-*TiAP1* vector. The resulting plasmid pMUbi-*TiAP1* and the vector harboring the *Bar* gene were mixed and co-bombarded with a particle bombardment of 1100 psi, 1 μg per gun, and 60 μg gold powder into the callus that was induced from the mature embryos of the Bobwhite wheat, which was highly susceptible to *Bgt*. During the induction and regeneration, the transformed wheat was screened using the herbicide Basta, while the presence of the transgenes was determined by amplifying the target gene using the pUbi-ASPF/R primers (Table S6).

Next, the disease responses to powdery mildew of the transgenic plants were tested. When the Bobwhite and the transgenic Bobwhite with the *TiAP1* gene grew to the three-leaf stage, the first leaves of them were spread out on an acrylic plate with the conidium of the *Bgt* race *E09* sprayed evenly. The leaves were harvested at 48 h post-inoculation with *E09* and stained with Coomassie Brilliant Blue as described previously (Göllner *et al*., 2008) and three biological replicates were conducted per stains with three technical replicates. Furthermore, to evaluate the resistance to *Bgt* of the Bobwhite and the transgenic Bobwhite with the *TiAP1* gene in the adult stage, they were natural infected by powdery mildew, where the infection type (IT) of each plant was scored as described by Lu *et al*. (2020).

### DNA and RNA extraction and qRT-PCR

Total DNA was extracted from the plant leaves using the cetyltrimethylammonium bromide (CTAB) method (Allen *et al*., 2006). While total RNA (500 ng) was extracted using the EasyPure^®^ Plant RNA kit (TransGen Biotech Co., Ltd, Beijing, China), reverse-transcribed (RT) using a RevertAid First Strand cDNA synthesis kit (TransGen Biotech Co. Ltd, Beijing, China), and quantified on an ABI Quantitative PCR Q6 Detection System (Thermo Fisher Scientific, USA) with the SYBR Premix Ex Taq kit (TransGene Biotech Co., Ltd, Beijing, China). The wheat actin gene was used as the reference gene, in which three independent biological replicates were conducted per analysis with at least three technical replicates. The primers have been listed in Table S6.

### Subcellular localisation

The CDS of the *TiAP1* fragments was fused green fluorescent protein (GFP) in the vector pCAMBIA1300. Then the constructed TiAP1-GFP or Δ without signal peptide of TiAP1 respectively) were transferred into the *Agrobacterium tumefaciens* GV3101 and transiently expressed in *N. benthamiana* leaves. Two days after infiltration, they were observed using a confocal laser scanning microscope (Leica SP5 X, Germany), Ten *N. benthamiana* leaves were analyzed in each of the three experiments and the detailed method was described in Methods S1.

### Transient expression of the TiAP1 protein induced by *Bgh* using barley leaves

The expression vector of the TiAP1 protein was constructed using the gateway system (Thermo Fisher Scientific, USA). The two recombinant plasmids, TiAP1-mYFP and mCherry-TaSYP51, were co-bombarded onto the leaves of the 7-day-old barley by particle bombardment at a helium pressure of 1,100 psi and placed on the 1% Phytagel^TM^ petri dishes containing 40mg ml^-1^ of benzimidazole (Sigma-Aldrich, USA) to culture for 24 h, and then inoculated with *Bgh* spores. After additional 24 h of inoculation, the expression of TiAP1 and TaSYP51 proteins were observed and analysed using a confocal laser-scanning microscope (Leica SP5 - X, Germany) according to the methodology of Smigielski *et al*. (2019). And three biological replicates were conducted per co-bombarded with six leaves.

### Protein Interaction Assay

We sampled the leaves of YN15 and SN6306, which were infected with *Bgt* from 0 to 5 d, and then construct a primary cDNA library using the BP reaction of the CloneMiner II kit (Invitrogen, Thermo Fisher Scientific, USA) by Qingdao Oebiotech Co. Ltd. (China). The quality of the cDNA library was identified. Next, we used the nuclear system Y2H mating standard operating procedure method as described in the manufacturer’s instructions (Clontech Laboratories, Inc., USA) to screen the protein that interacts with TiAP1 and further verified the obtained interaction protein, and three biological replicates were conducted per Y2H verify. The structure, amino acid sequence domain, and the signal peptide of the obtained protein BgCDA1 were analysed using the the Basic Local Alignment Search Tool (PBLAST; Altschul *et al*., 1997), Pfam (Finn *et al*., 2014), and SignalP 5.0 (Almagro Armenteros *et al*., 2019) software, respectively.

For LCI assay, by amplifying the sequence of TiAP1 and BgtCDA1 and applying homologous recombination with the nLuc and cLuc, as previously described (Chen *et al*., 2008), the fusion protein and one of the corresponding co-injected empty vectors, nLuc/cLuc, was co-expressed in *N. benthamiana* as a negative control. At 48 h post-infiltration, the LUC activity of the leaves was observed using an *in vivo* imaging system (Berthold LB985 NightSHADE, Germany). Ten *N. benthamiana* leaves were analyzed in each of the three experiments.

For BiFC assay, the sequences of TiAP1 and BgtCDA1 were amplified and fused with the N and C terminals of the GFP, respectively, using the gateway system (Thermo Fisher Scientific, USA). As mentioned previously (Bracha-Drori *et al*., 2004), the fusion protein and one of the corresponding empty GFP-N/GFP-C vectors were co-expressed in *N. benthamiana* as a negative control. The GFP signal was observed 48 h after infiltration using a confocal laser-scanning microscope (Leica SP5-X, Germany). Ten *N. benthamiana* leaves were analyzed in each of the three experiments.

### The yeast expression, purification and chitin deacetylase activity assay of BgtCDA1

For protein expression and purification in yeast, BgtCDA1 coding sequences with a 6XHIS tag were amplified and ligated into pPIC9K-HIS vector (Invitrogen), and then transformated into *Pichia pastoris* strain GS115 (Invitrogen). The positive transformants were induced protein expression using methanol. The target proteins were purified using Ni-column affinity chromatography. Purified protein was used to detected the enzyme activity of BgtCDA1 by measuring the amount of released acetate using ion chromatography and qualitative analysis of deacetylated products by MALDI-TOF by Hoogen Biotech, Shanghai, China according to Gao et al., (2019). MALDI-TOF MS analysis was performed on a Ultraflex Extreme MALDI-TOF-TOF mass spectrometer (Bruker Daltonics, USA). The standard reaction mixture (200 μl) μM BgtCDA1, 50 mM Tris-HCl (pH8.0) and 1 mM chitin oligomers with 6 GlcNAc moieties (A6) as the substrate were incubated at 37 ℃ for 5, 30 min followed by heating at 100 ℃ for 10 min. The mixture without BgtCDA1 was as the control. The yeast expression, purification and enzyme activity assay of BgtCDA1 were detailed performed described in Methods S1.

### RNA-Seq and data analysis

RNA-Seq analysis was performed using the two-leaf stage Bobwhite (Bob-0) and the transgenic Bobwhite with the *TiAP1* gene (OE2-0) as well as a Bobwhite inoculated with *E09* two days later (Bob-2) and the corresponding transgenic Bobwhite with the *TiAP1* gene (OE2-2). Three biological replicates were used for the RNA extraction, where total RNA was extracted and treated with DNase I, in which the quality test was performed, and then used to construct the library. The quality of the library was assessed using an Agilent 2100 Bioanalyzer (Agilent Technologies, Inc., USA), while the libraries were sequenced on an Illumina HiSeq 2500 platform (Illumina, Inc., USA). The RNA-Seq reads were aligned to the wheat genome of the International Wheat Genome Sequencing Consortium with the RefSeq v1.1 annotation (https://wheat-urgi.versailles.inra.fr/Seq-Repository/Annotations). Differentially expressed genes across the samples were identified using the DESeq2 package (Love *et al*., 2014) using the standard parameters with an adjusted false-discovery rate of *P*-value < 0.01 and fold change > 4.

### Statistical analysis

The statistical significance of the results was calculated using a one-way analysis of variance followed by least significant difference and Duncan’s new multiple range test at a significant difference (*P* < 0.05) using IBM SPSS software version 19.0 (Chicago, Illinois, USA). All experiments reported include three to six biological replicates and a minimum of three technical replicates with similar results. All primers have been listed in Table S6.

## Accession numbers

The cDNA and coded protein sequence of *TiAP1* sequence in this research has been deposited in NCBI under the accession number KJ513672 and AJC64141.1; *Blumeria graminis* f. sp. *tritici* BgtCDA1 was coded_by =” join (KE374986.1:122515..122759, KE374986.1:122809..123142, KE374986.1:123188..123296, KE374986.1:123342..123381, KE374986.1:123432..123502, KE374986.1:123547..123801, KE374986.1:123919..123923, KE374986.1:123978..124142, KE374986.1:124319..124375), and had one and two additional alternative splicings compared with EPQ66796 between KE374986.1:123143 and KE374986.1:123187, KE374986.1:123802 and KE374986.1:123977, respectively; Other protein sequence data from this article can be found in the NCBI under the following accession numbers: *Saccharomyces cerevisiae CDA1 (NP_013410) and Saccharomyces cerevisiae CDA2 (NP_013411).* TaSYP51 was coded by sequence in chr5A of wheat between 67321475 and 67319313.

## Acknowledgements

This work was supported by the grants from the National Natural Science Foundation of China (31771777); the Key R & D program of Shandong Province (Public Welfare Special) (2018GSF121008); Shandong ‘Double Tops’ Program; the Major Basic Research Project of Shandong Natural Science Foundation (2017C03); and the Overseas Visiting Programme for Graduate Mentors of Shandong Province. We are very grateful to Prof. Hans Thordal-Christensen’s group, University of Copenhagen, Denmark, for providing experimental assistance, to Dr. Patrick Schweizer, IPK, Gatersleben, Germany, for supplying the RNAi vector and the Mlo-RNAi construct, to Prof. Pinghua Li, Shandong Agricultural University, for providing the LCI system, and to Prof. Dawei Li, China Agriculture University, for providing the BSMV-RNAi system. Confocal image data were collected at the Center for Advanced Bioimaging (CAB), University of Copenhagen, Denmark.

## Author contributions

D.F., H.W., and Y.Y. conceived and designed the experiments. Y.Y., P.F., J.L., W.X., N.L., Z.N., Q.L., J.S., Q.T., and Y.B. performed the experiments and analysed the data; Y.Y. and D.F. wrote the paper; D.F. and H.W. conceived, directed and coordinated the project.

## Supporting Informations Short Legends

Fig. S1 The TiAP1 prokaryotic expression and purification. (A) The TiAP1 protein expression was detected by SDS-PAGE. M: Low-molecular weight protein ladder, Lane1: Uninduced TiAP1; Lane 2: 15 °C induced overnight. (B) TiAP1 quality assurance after renaturation. M: Low-molecular weight protein ladder; Lane 1: Bovine serum albumin (2.0 µg); Lane 2: TiAP1 protein (1.5 µg).

Fig. S2 Identification of the transgenic Bobwhite with the *TiAP1* gene and the subsequent hybrid generation. (A) Identification of the *Bar* gene in the transgenic *TiAP1* wheat. M: DL2000 DNA Marker [Takara Biomedical Technology (Beijing) Co., Ltd.]; Lane 1-10: transgenic lines. (B) Identification of the *TiAP1* gene in transgenic wheat. M: Trans2K^®^ Plus II DNA Marker [TransGen Biotechnology (Beijing) Co., Ltd.]; Lane 1-2: Bobwhite and ddH_2_O; Lane 3-12: transgenic lines. (C) Identification of the *TiAP1* gene in the subsequent hybrid generation of the wheat F_3_. M: DL2000 DNA Marker [Takara Biomedical Technology (Beijing) Co., Ltd.]; Lane 1-10: subsequent hybrid generation of the F_3_ lines; Lane 11: OE1; Lane 12: YN15 wheat.

Fig. S3 Validation of wheat powdery mildew resistance in the representative hybrid subsequent generation of F_3_, which showed resistance to *E09* at 7 dpi in the seedling stage. Wheat YN15 and OE1 were used as the susceptible and resistant controls, respectively. Lane 1: YN15; Lane 2-11: The subsequent hybrid generation of the F_3_ lines 1-10; Lane 12: OE1.

Fig. S4 Validation of the wheat powdery mildew resistance in 7 transgenic wheats Bobwhite with the *TiAP1* gene in the adult stage, which showed high resistance to powdery mildew. Bobwhite was used as the susceptible control. (A) Disease phenotype of Bobwhite and the transgenic Bobwhite with *TiAP1* when inoculated with *E09* for ten days. (B) The proportion of lesions in the leaf area of Bobwhite and the transgenic Bobwhite with *TiAP1* when inoculated with *E09* for ten days.

Fig. S5 Subcellular localisation of the fusion protein in the *N. benthamiana* leaf epidermal cells. (A) GFP, TiAP1-GFP and Δsp-TiAP1-GFP under normal conditions for GFP signal detection in *N. benthamiana* leaves, where GFP acted as a positive control signalling the nucleus and cytoplasm, while Δsp-TiAP1-GFP was localised in the cytoplasm. (B) Western blot detection of the GFP, TiAP1-GFP and Δsp-TiAP1-GFP proteins expression.

Fig. S6 The BgtCDA1 protein bioinformatics analysis. (A) The domain of the amino acid sequence of BgtCDA1 by Pfam; (B) The structure analysis of BgtCDA1 by the Basic Local Alignment Search Tool (BLAST); (C) The SignalP analysis of the amino acid sequence of BgtCDA1. (D) Alignment of the five conserved motifs of the BgtCDA1 domain of *Saccharomyces cerevisiae* CDA1 (NP_013410) and *Saccharomyces cerevisiae* CDA2 (NP_013411) as well as the five conserved motifs (Motif 1-5) of the carbohydrate esterase family 4 (CE-4) that are indicated by red boxes.

Fig. S7 Transcriptome analysis of the *TiAP1* and *BgtCDA1* co-regulated genes. (A) Sample-to-sample cluster analysis results. (B) The pie chart shows the different expression and up-regulated genes in OE2-2 relative to Bob-2. (C) Heat map showing the differentially expressed genes (DEGs) inoculation with *E09* for two days. (D) Gene ontology (GO) analysis of the top 30 DEGs in OE2-2 and Bob-2.

Methods S1 Additional description of methods.

Table S1 Evaluation of the resistance to *E09* in YN15, wheat lines overexpressing *TiAP1*, and the hybrid subsequent generation of F_3_ in the seedling stage. Lines 1-10: ten hybrid F_3_ generation lines. YN15 and OE1 are as the susceptible and resistant controls respectively

Table S2 Evaluation of the resistance to powdery mildew in Bobwhite and wheat lines overexpressing *TiAP1* at the adult stage. OE 1-10 represent ten wheat lines overexpressing *TiAP1*. Bobwhite is as the control

Table S3 Differential gene expression in the transgenic lines overexpression OE2 and Bobwhite when inoculated with *E09* for two days. The default conditions of the screening included *P* < 0.01 and fold change > 4

Table S4 Gene ontology (GO) analysis of the differentially expressed genes in OE2-2 and Bob-2

Table S5 Expression of selected pathogen-responsive genes in Bob-0, Bob-2, OE2-0, and OE2-2 in the RNA-Seq experiments

Table S6 The primers used in this study

Video S1 Confocal microscopy (Leica SP5-X, Germany) z-scan image series of TiAP-mYFP/mCherry-TaSYP51 on barley leaf epidermic cell under the infection of *Bgh*. Bar = 20 µm.

