## Supplementary figures, tables, methods, and movie and for "*Thinopyrum intermedium* TiAP1 interacts with a chitin deacetylase from *Blumeria graminis* f. sp. *tritici* and increases the resistance to *Bgt* in wheat": Methods S1.docx

**Supplementary methods**

**Virus-induced gene silencing (VIGS)**

VIGS was conducted according to Yuan *et al*. (2011) to insert the conserved sequence of the target *TiAP1* gene into the γ vector and transform the *Agrobacterium*. The three types of *Agrobacteria* harbouring α, β, and BSMV-TiAP1as or BSMV-BgtCDA1as, respectively, were resuspended in an infiltration buffer that was adjusted to an optical density (OD)_600_ of 0.7 and incubated at 25 °C for 3 h. Then mixed at a ratio of 1:1:1, to inject into the leaves of *N. benthamiana* at the 4-8 leaf stage. After being maintained in a growth chamber for 12 days post-infiltration (dpi), the infiltrated leaves were harvested and grind in 20 mM Na-phosphate buffer (pH 7.2) containing 1% celite, where the sap was mechanically inoculated onto the leaves of the two-leaf stage trititrigia SN6306 or YN15.

**Host-induced gene silencing (HIGS)**

The conserved sequence of *BgtCDA1* gene was obtained by a multiple sequence alignment, referring to the method of Nowara *et al*. (2010) and using the SI-FI software (http://labtools.ipk-gatersleben.de/) to predict the RNA interference (RNAi) off-target genes in the *Bgt* genome database (https://www.ncbi.nlm.nih.gov/Taxonomy/Browser/wwwtax.cgi?lvl=0&amp;id=1268274). The conservative region of *BgtCDA1* was recombined with the pIPKTA30N RNAi vector, where we applied a particle bombardment to co-transform the HIGS plasmid, BgtCDA1 RNAi, and a β-glucuronidase (GUS) reporter gene construct into the leaf epidermal cells of the wheat YN15 that grew for seven days. Seven leaves were transformed per construct. Two days later, the leaves were inoculated with *Bgt* and three days later, the leaves were stained for GUS activity. The fungal haustorial formation was examined for the transformed (GUS expressing) blue cells using a biological microscope (Nicon Ni-U). The haustorial index (HI) was computed as the ratio of transformed cells with haustoria divided by the total number of transformed cells. The empty vectors pIPKTA30N, and the Mlo-RNAi (pIPKTA36) constructs, were used as negative and positive controls, respectively, while the HI was calculated relative to the empty vector in each experiment, which was set to 100%, according to Ahmed (2015).

**The TiAP1 protein purification**

The entire bacterial sample was added to a lysis buffer [50 mM Tris (pH 7.5) with 150 mM NaCl containing 1% Triton X-100, 1 ug/mL pepstatin A, and 1 ug/mL leupeptin], where the mixture was disrupted by ultrasonic disintegration in an ice-water bath. At the same time, the nickel iminodiacetic acid (Ni-IDA) affinity column was balanced with 50 mM Tris (pH 7.5) and a 150 mM NaCl buffer. The inclusion bodies were rinsed with a wash buffer [50 mM Tris (pH 7.5) with 150 mM NaCl, containing 1% Triton X-100, 5 mM ethylenediaminetetraacetic acid (EDTA) and 2 mM dithiothreitol (DTT)], followed by a 50 mM Tris (pH7.5), 150 mM NaCl, and 8 M urea buffer that dissolved the inclusion bodies and balanced the Ni- IDA column. Finally, the TiAP1 protein was eluted with a 50, 100, and 300 mM imidazole elution buffer, where each eluted fraction was collected for SDS-PAGE analysis and detection.

The fractions containing the TiAP1 fusion protein with the 300 mM imidazole were pooled and extensively dialysed against the digestion buffer [50 mM Tris (pH 7.5), 150 mM NaCl, 2 mM EDTA, 4 mM glutathione (GSH), 0.4 mM oxidized glutathione (GSSG), 0.4 M L-arginine, and 2 mM DTT] at 4 °C for protein renaturation. The recombinant TiAP1 protein was then dialysed in the storage solution (50 mM Tris, 150 mM NaCl, 2 mM DTT, 10% glycerol, pH 7.5). After the dialysis, the supernatant was filtered with a 0.22 µm filter, aliquoted, and frozen to –80 °C, where the label protein expressed by the pET28a vector was purified as described earlier and used as a control.

**Transient Expression in Tobacco and Observation of GFP**

The subcellular localization expression vector was constructed using the CDS of the *TiAP1* fragments that were cloned into the *Xba* I - *Asc* I sites of the digested pCAMBIA1300 to construct the pCambia1300-*TiAP1* vector. Then the constructed TiAP1-GFP, Δsp-TiAP1-GFP (with and without signal peptide of TiAP1 respectively) and GFP were transferred to the *Agrobacterium* *tumefaciens* GV3101. The free GFP was as control. The cultured cells were resuspended in buffer [10 mM MgCl_2_, 10 mM 2-(N-morpholino) ethanesulfonic acid (MES), pH 5.7, and 0.1 mM acetosyringone (AS)], where the OD_600_ was adjusted to 0.5-0.6, placed in the dark at 25 °C for 3 h, and then infiltrated into *N. benthamiana* leaves using a needleless syringe. A confocal laser scanning microscope (Leica SP5‐X) under the excitation of 488 nm and emission of 507 nm were used to observe the fluorescence, while plasmolysis was obtained by injecting 0.8% mannitol.

Proteins separated by SDS-PAGE were transferred to an 0.22 um polyvinylidene difluoride (PVDF) membrane (pretreated with methanol for 30 s) using the Mini Trans-Blot apparatus (Bio-Rad) at 400 mA for 40 min. Equal protein transfer was monitored by staining the membranes with Ponceau S (Sigma–Aldrich). To reduce non-specific binding, the membrane can be soaked in 10% skimmed milk (in PBS, pH 7.2) for 1 hour at room temperature, or overnight at 4°C. The following primary antibodies were used in this study: anti-GFP (diluted 1:3000, HT801-01, TransGen Biotech Co., Ltd, Beijing, China) and anti-Flag (diluted 1:3000, HT201-01, TransGen Biotech Co., Ltd, Beijing, China) antibodies. The secondary antibody goat anti-mouse *ProteinFind*^®^ Goat Anti-Mouse IgG (H+L), HRP Conjugate (diluted 1:3000, HS201-01, TransGen Biotech Co., Ltd, Beijing, China) was applied and proteins were detected using an ECL Western blotting detection system (FUSION FX SPECTRA, Vilber, France).

**The yeast expression, purification and chitin deacetylase activity assay of BgtCDA1**

For protein expression and purification in yeast, BgtCDA1 coding sequences with a 6XHIS tag were amplified and ligated into an *Eco*I-*Not*I-linearsized pPIC9K-HIS vector (Invitrogen). The construct was digested by *Sal*I, and then transformated into *Pichia pastoris* strain GS115 according to the manual of *Picchi* yeast expression (Invitrogen). The transformants were identified using the amplification by 5’AOX/3’AOX primer pairs and sequencing. The positive transformants were cultured in BMGY medium, and then transferred to BMMY medium to induce protein expression using methanol. The target proteins were purified using Ni-column affinity chromatography. Purified protein was examined by SDS-PAGE and Western blotting as previously described. The protein concentrations were determined with the protein assay kit (Bio-Rad) according to the instructions. The enzyme activity of BgtCDA1 was determined by measuring the amount of released acetate using ion chromatography and qualitative analysis of deacetylated products by MALDI-TOF by Hoogen Biotech, Shanghai, China according to Gao et al., (2019). MALDI-TOF MS analysis was performed on a Ultraflex Extreme MALDI-TOF-TOF mass spectrometer (Bruker Daltonics, USA). Mass spectra were obtained in the positive ion reflective mode. Reaction products (1ul) were spotted on the target plate, and dried following by 1 ul of 2,5-dihydroxybenzoic acid solution [20mg ml-1 in 50:50 acetonitrile/water containing 0.1% trifluoroacetic acid (TFA)] and dried following by MALDI-TOF MS analysis. The standard reaction mixture (200 μl) containing 2 μM BgtCDA1, 50 mM Tris-HCl (pH8.0) and 1 mM chitin oligomers with 6 GlcNAc moieties (A6) as the substrate were incubated at 37 ℃ for 5, 30 min followed by heating at 100 ℃ for 10 min. The mixture without BgtCDA1 was as the control.
