## Supplementary figures and images for "*Thinopyrum intermedium* TiAP1 interacts with a chitin deacetylase from *Blumeria graminis* f. sp. *tritici* and increases the resistance to *Bgt* in wheat"

### figure S1.tif

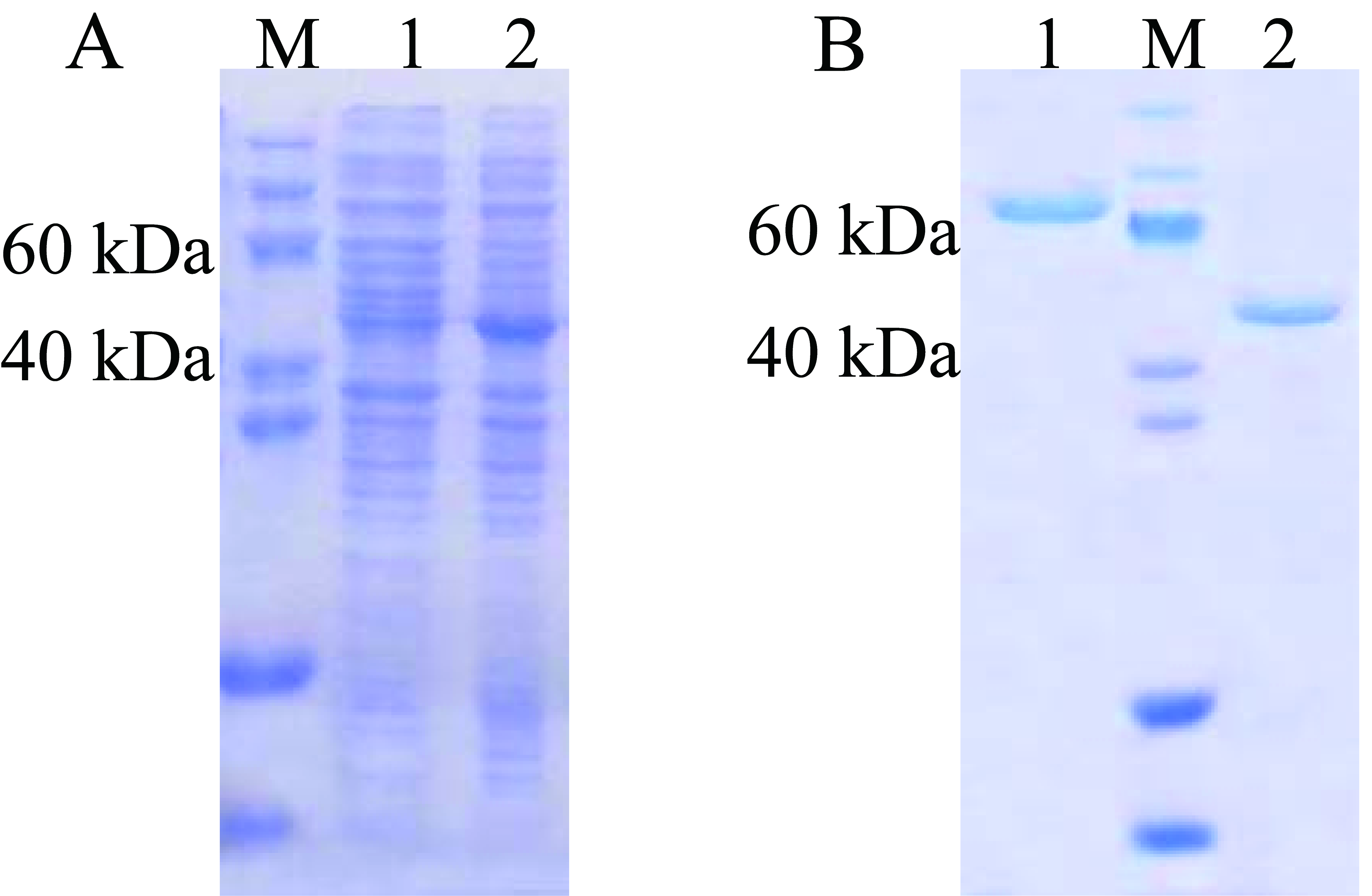

### figure S2.tif

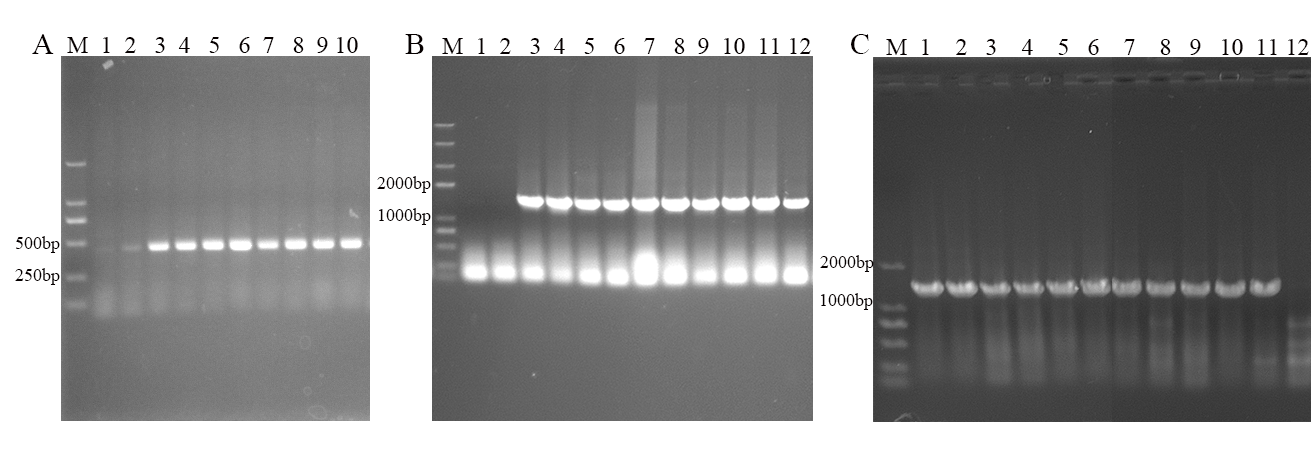

### figure S3.tif

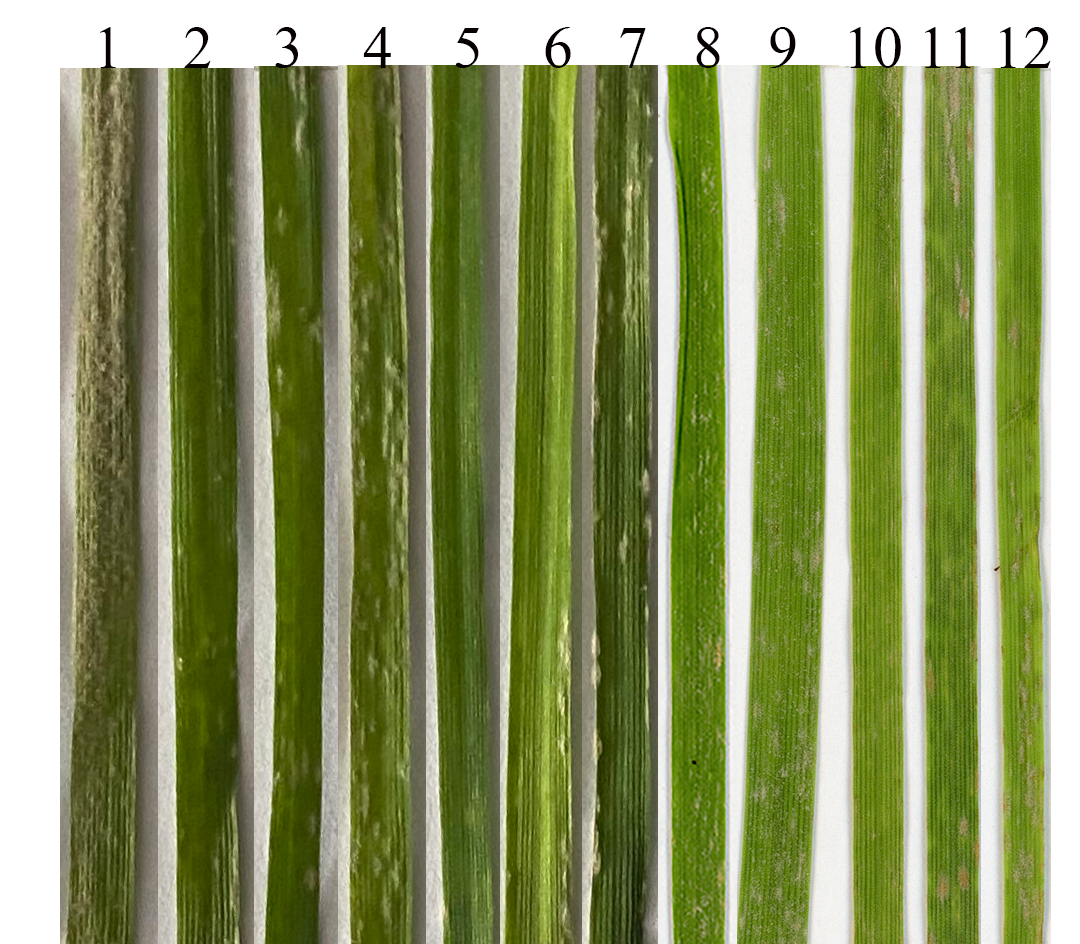

### figure S4.tif

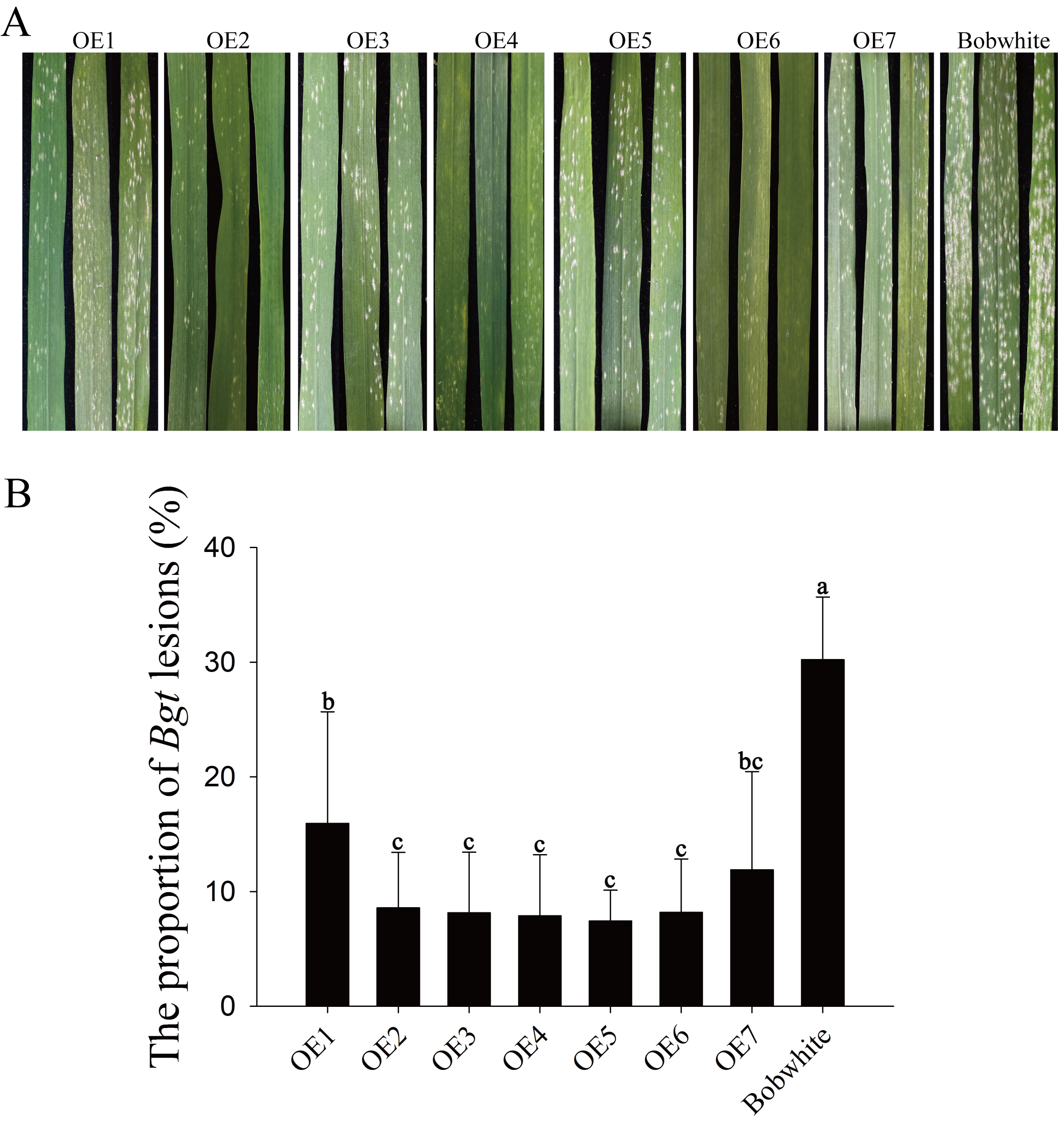

### figure S5.tif

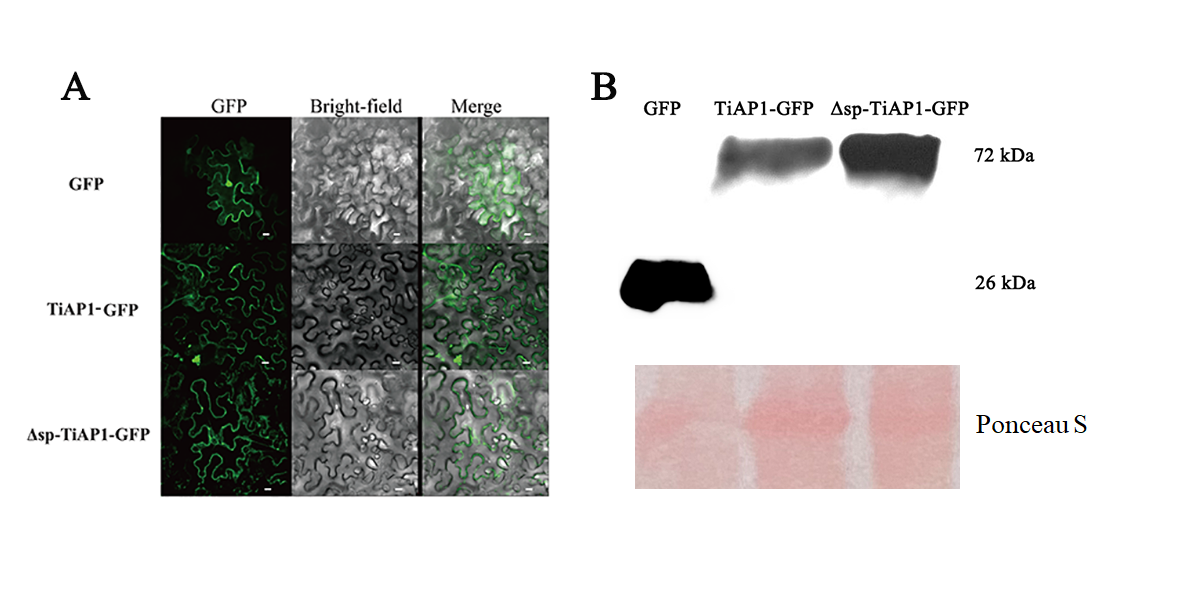

### figure S6 .tif

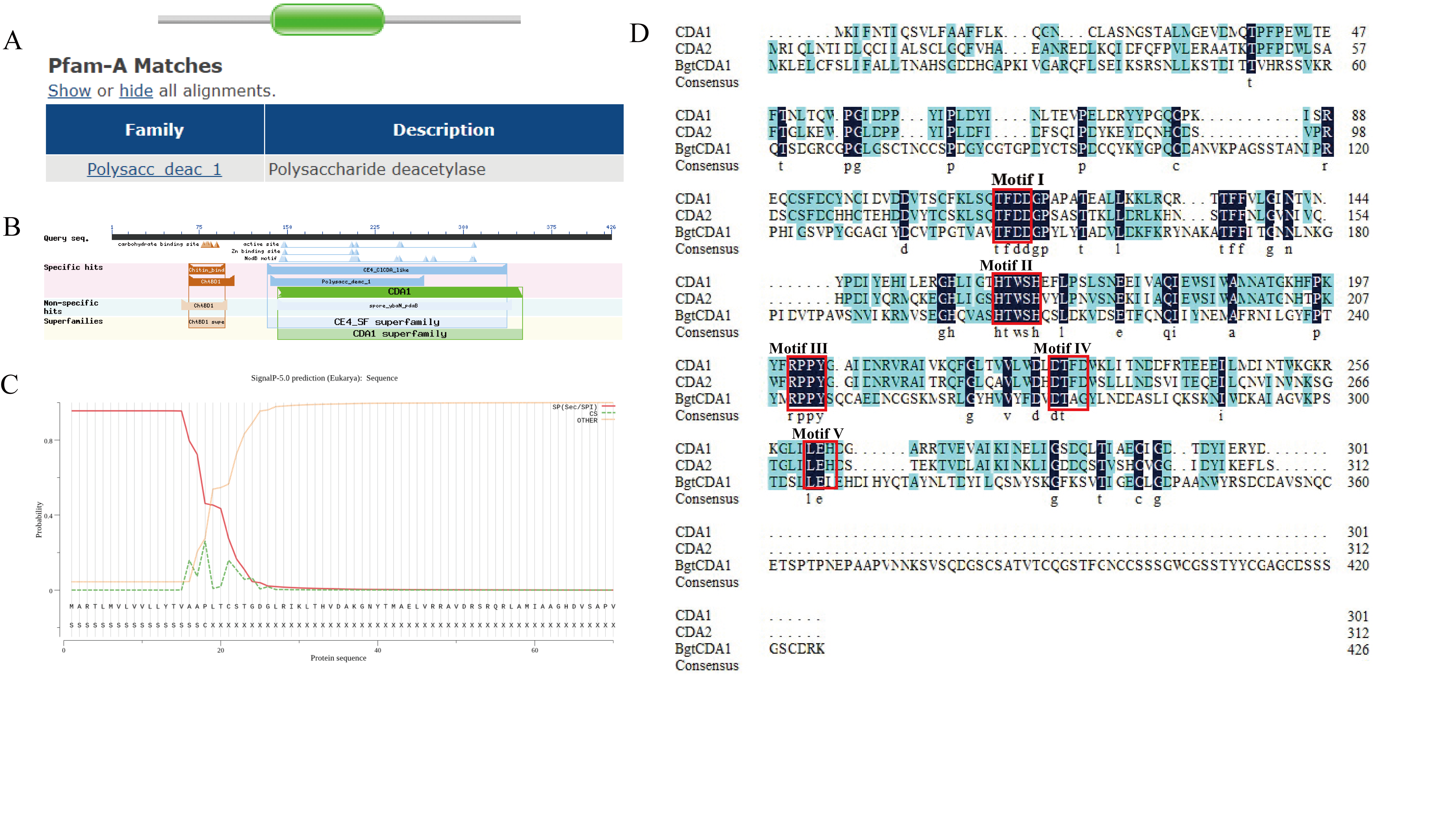
